# Multiplexed single-cell transcriptomics reveals diverse phenotypic outcomes for pathogenic SHP2 variants

**DOI:** 10.1101/2025.06.30.662374

**Authors:** Anne E. van Vlimmeren, Ross M. Giglio, Ziyuan Jiang, Minhee Lee, José L. McFaline-Figueroa, Neel H. Shah

## Abstract

The protein tyrosine phosphatase SHP2, encoded by *PTPN11*, is an important regulator of Ras/MAPK signaling that acts downstream of receptor tyrosine kinases and other transmembrane receptors. Germline *PTPN11* mutations cause developmental disorders such as Noonan Syndrome, whereas somatic mutations drive various cancers. While many pathogenic mutations enhance SHP2 catalytic activity, others are inactivating or affect protein interactions, confounding our understanding of SHP2-driven disease. Here, we combine single-cell transcriptional profiling of cells expressing clinically diverse SHP2 variants with protein biochemistry, structural analysis, and cell biology to explain how pathogenic mutations dysregulate signaling. Our analyses reveal that loss of catalytic activity does not phenocopy SHP2 knock-out at the gene expression level, that some mechanistically distinct mutations have convergent phenotypic effects, and that different mutations at the same hotspot residue can yield divergent cell states. These findings provide a framework for understanding the connection between SHP2 structural perturbations, cellular outcomes, and human diseases.

## Introduction

SHP2, encoded by *PTPN11*, is a ubiquitously expressed protein tyrosine phosphatase in humans that functions as a signaling hub downstream of many transmembrane receptors and has critical roles in cell proliferation, cell differentiation, immunity, and development (**Figure 1A**). *PTPN11* missense mutations drive many human diseases, including hematopoietic malignancies such as acute myeloid leukemia (AML), acute lymphoid leukemia (ALL), and juvenile myelomonocytic leukemia (JMML) (**Figure 1B**). Association with solid tumors such as neuroblastoma, hepatocellular carcinoma, glioblastoma, and melanoma has also been described^1–3^. Germline mutations in *PTPN11* underlie congenital disorders, including approximately 50% of cases of Noonan Syndrome (NS) cases^4^, and 95% of Noonan Syndrome with Multiple Lentigines (NSML) cases^5,6^ (**Figure 1B**). NS is characterized by facial dysmorphia, intellectual disability, and heart defects – in particular pulmonic stenosis^4^. In addition to these NS-phenotypes, NSML patients have a high incidence of hypertrophic cardiomyopathy, electrocardiographic abnormalities, and hearing loss^6^.

**Figure 1.**
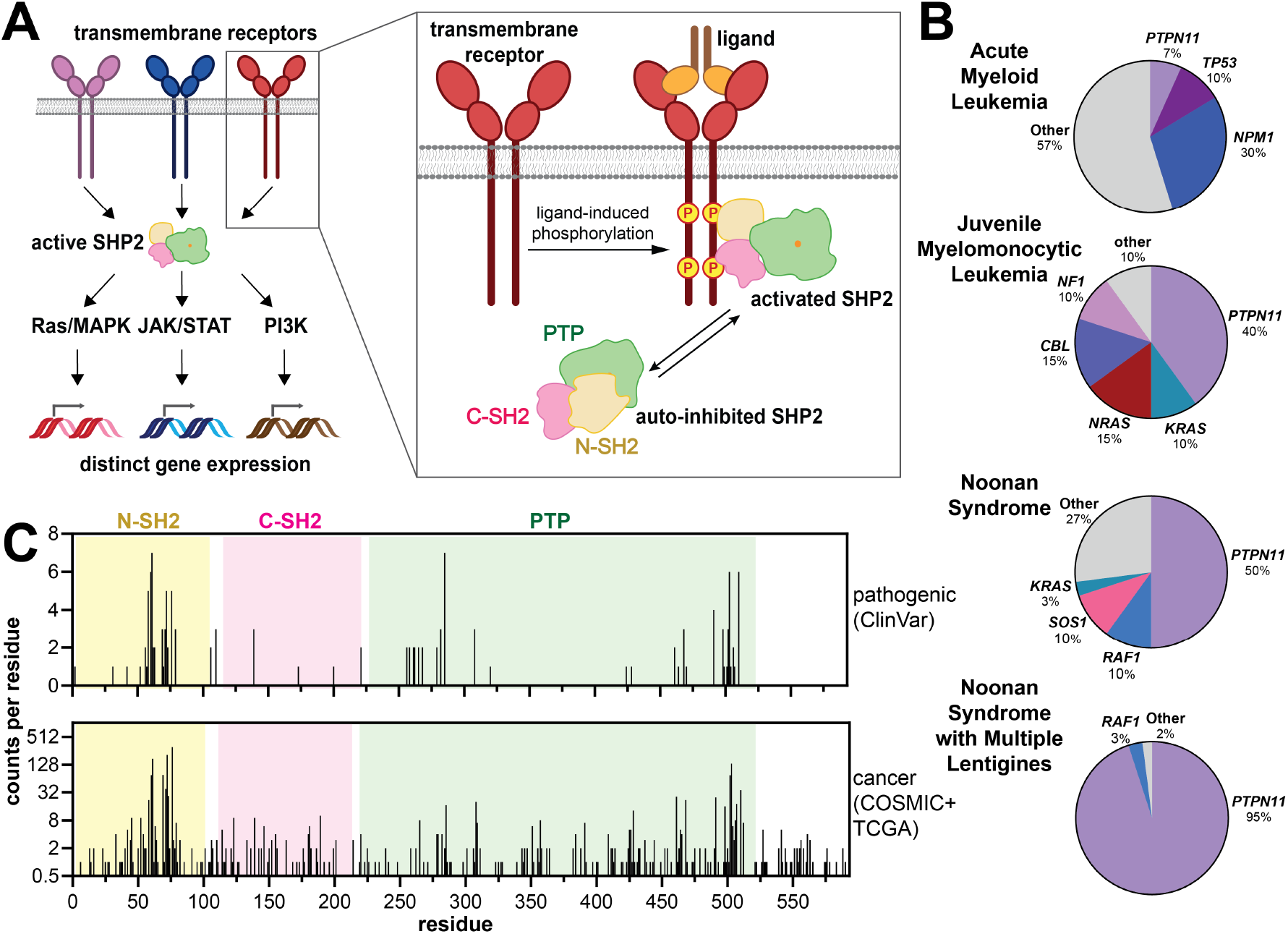
Biological function and pathology of SHP2. (**A**) SHP2 receives input from a variety of cell signaling pathways, and SHP2 activation by binding to phosphoproteins has a diverse array of potential signaling and transcriptional outcomes. (**B**) Pie charts showing disease-driving genes for various human diseases, as identified in DNA sequencing of patient cohorts^4,28–33^. *PTPN11* mutations underlie both congenital disorders and cancers. (**C**) Positions and frequencies of missense mutations in SHP2 along its 593-residue sequence. Pathogenic mutations were obtained from the ClinVar dataset. Cancer-associated mutations were obtained from the COSMIC and TCGA databases.

Most disease-associated functions of SHP2 have been attributed to its role in Ras/MAPK signaling. Indeed, both NS and NSML are categorized as “RASopathies” and share similarities with other syndromes caused by mutations in Ras/MAPK components. In many cancers, SHP2 mediates signal transduction from receptor tyrosine kinases to Ras, and several allosteric inhibitors of SHP2 have entered clinical trials for the treatment of receptor tyrosine kinase-driven cancers^7,8^. SHP2 promotes Ras/MAPK signaling through several mechanisms, including inhibition of Sprouty1, a negative regulator of Ras^9^, direct dephosphorylation and activation of Ras, and dephosphorylation of scaffold proteins to prevent the recruitment of Ras GTPase activating-proteins (RasGAPs) to signaling complexes^10–12^. It is noteworthy, however, that SHP2 functions downstream of a variety of transmembrane receptors and can activate not just the Ras/MAPK pathway, but also PI3 kinase signaling, JAK/STAT signaling, and immune checkpoint signaling (**Figure 1A**)^13–16^.

Hundreds of SHP2 mutations are cataloged in clinical databases (**Figure 1C**), and these mutations disrupt SHP2 structure and function through diverse mechanisms^17–21^. SHP2 is canonically activated by binding to phosphoproteins, which disrupts its resting auto-inhibited state to yield an active enzyme (**Figure 1A**) ^22,23^. Many oncogenic mutations also disrupt auto-inhibition, leading to catalytic gain-of-function effects^17,23^. By contrast, other mutations appear to act through non-catalytic mechanisms^24^. Despite extensive studies, how these molecular effects translate into disease phenotypes remains unclear. One emerging theme is that SHP2 mutations can alter protein-protein interactions, as seen in NSML-associated variants. Many NSML mutations result in low or no catalytic activity but also cause large conformational changes that enhance binding to MPZL1/Pzr, a driver of hypertrophic cardiomyopathy through the Akt and NF-κB pathways^25^. Our recent work suggests that many pathogenic mutations in SHP2 broadly reshape its protein interaction network, yet the transcriptional and signaling consequences of these changes remain poorly understood^19,26^.

Subtle perturbations to the structure of a signaling protein, such as those caused by missense mutations, can propagate to changes in protein-protein interactions and proximal signaling events, which in turn, can alter downstream gene expression. Indeed, a previous study on the transcriptomes of cells expressing two SHP2 mutants that disrupt auto-inhibition found increased expression of metabolic proteins, highlighting the potential insights that could be gained from studying mutation-specific transcriptional changes^27^. In addition, profiling mutation-driven changes in gene expression may also uncover key disruptions to protein function and can be leveraged to aid our understanding of SHP2 structure-function relationships. Here, we use single-nucleus RNA sequencing to map the transcriptional impact of 15 clinically and mechanistically diverse pathogenic *PTPN11* mutations. We identify SHP2 presence as a critical driver for the cellular response to epidermal growth factor (EGF) stimulation and demonstrate that the R138Q mutation, which prevents C-SH2 domain interactions, attenuates EGF-driven signaling independent of catalytic activity. Further, we show that two mutations, T507K and Q510K, have convergent effects on the biochemical and transcriptional level, with charge of the resulting amino acid as a likely driver. Finally, we show that different disease-relevant substitutions at catalytic residue Q510 have different effects on SHP2 structure and activity, propagating to distinct transcriptional outcomes. By systematically profiling the transcriptional landscape of *PTPN11* variants, we provide new insights into how SHP2 mutations alter protein function.

## Results

### Single-cell transcriptomics identifies global transcriptional changes induced by SHP2 expression

To profile the effects of a collection of pathogenic SHP2 mutations on gene expression networks alone and under mitogen stimulation, we used sci-Plex-v2 multiplex single-cell RNA sequencing^34,35^. We transfected either SHP2^WT^ or mutant SHP2 into a SHP2 knock-out (SHP2^KO^) HEK 293 cell line **(Figure 2A,B and Supplementary Figure 1A**). Cells were stimulated with a range of EGF concentrations or left unstimulated, then nuclei for each condition were harvested 24 or 96 hours post-stimulation and uniquely barcoded by fixation of an oligonucleotide hash. Barcoded nuclei were pooled, cDNA processed, and single-nuclei mRNA libraries were generated using our modified version of combinatorial indexing RNA-seq^34–38^. We captured a total of 29,716 cells across two replicates with a mean of 2447 cells per SHP2 variant and a mean coverage of 155 cells per unique combination of SHP2 variant, EGF dose, and time point (**Supplementary Figure 1B-D**).

**Figure 2.**
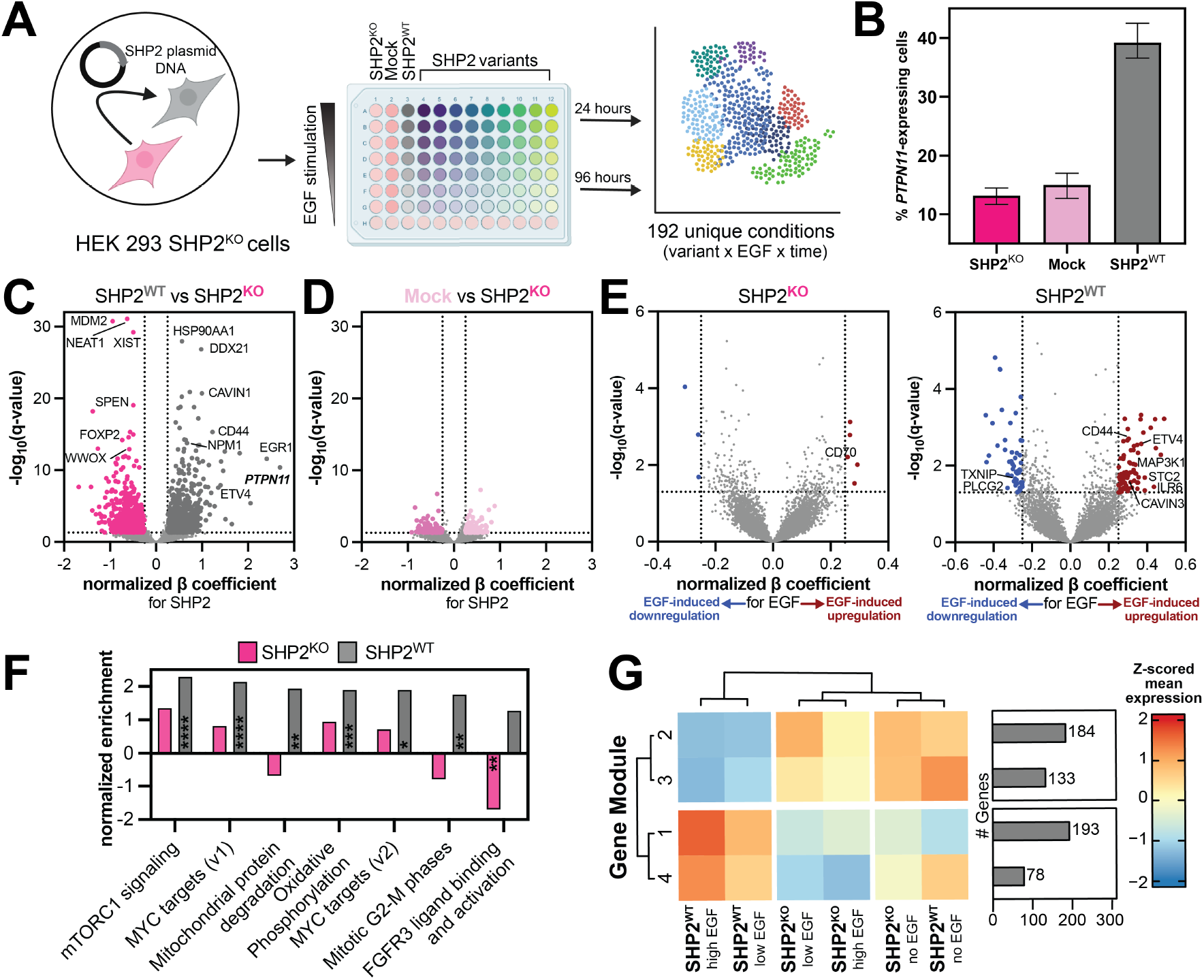
Single-nucleus RNA-sequencing reveals the transcriptional effect of SHP2 expression. (**A**) Schematic overview of RNA-sequencing experiment to probe effects of SHP2 on gene expression. (**B**) Percentage of *PTPN11*-expressing cells in the SHP2^KO^ population, SHP2^WT^-transfected cells, and mock-transfected cells, out of total number of cells sequenced for those respective samples. (**C**) Volcano plots showing SHP2-induced differentially expressed genes (DEGs) for SHP2^KO^ and SHP2^WT^ at 24 hours. Any significant (false discovery rate < 0.05) transcript with a β coefficient of > 0.25 or <-0.25 is colored. (**D**) Same as (**C**), but for mock-transfected vs SHP2^KO^. (**E**) Volcano plots showing EGF-induced differentially expressed genes for SHP2^KO^ (*top*) and SHP2^WT^ (*bottom*). Any significant (false discovery rate < 0.05) transcript with a normalized β coefficient of > 0.25 or <-0.25 is colored. (**F**) Gene Set Enrichment Analysis shows pathways of DEGs for SHP2^WT^ and SHP2^KO^. * denotes false discovery rate <0.05, ** < 0.01, *** < 0.001, and **** <0.0001. (**G**) Four gene modules were identified between SHP2^WT^ and SHP2^KO^. SHP2^WT^ without EGF stimulation behaves most similar to SHP2^KO^. Low EGF is defined as 12.5-50 ng/mL; high EGF is defined as 100-1000 ng/mL).

First, we established the effect of SHP2^WT^ presence on gene expression, by comparing SHP2^KO^ and SHP2^WT^ cells across EGF-stimulation conditions (**Figure 2C and Supplementary Table 1**). We juxtaposed this with a comparison between SHP2^KO^ and mock-transfected cells, as our control (**Figure 2D**). Between mock-transfected cells and SHP2^KO^ cells, we identified 114 genes as significantly upregulated (quasipoisson regression, > 0.25 β coefficient, < 0.05 false discovery rate (FDR)) in mock-transfected cells, and 105 genes that were downregulated (< -0.25 β coefficient, < 0.05 FDR) (**Figure 2D and Supplementary Figure 1E**). By contrast, we identified 820 genes that were significantly upregulated in cells expressing SHP2^WT^, including *PTPN11* itself (**Figure 2C and Supplementary Figure 1E**). These genes were enriched for Gene Ontology (GO) terms related to kinase signaling and cell cycle, as well as nuclear export and mitochondrial import (**Supplementary Figure 2A and Supplementary Table 1**). Moreover, several known EGF-response genes were also enriched, including *EGR1/3, ETV4/5, JUN, ATF5, CCND1, and DUSP1*^39^, indicating that the mere presence of SHP2 promotes the cellular response to EGF-stimulation (**Figure 2C and Supplementary Figure 2B**). This is consistent with the observation that receptor tyrosine kinase-driven cancer cell lines depend on SHP2 for proliferation^40^. Interestingly, these genes remained highly expressed in SHP2^WT^ cells compared to SHP2^KO^ at 96 hours (**Supplementary Figure 2C,D**). 738 genes were downregulated in SHP2^WT^-expressing cells compared to SHP2^KO^ (**Figure 2C and Supplementary Table 1**). Notably, we detected altered expression of genes related to heart development, mesenchymal stem cell (MSC) differentiation, and telencephalon development in SHP2^KO^ cells, aligning with the known roles of SHP2 in cardiac pathology, MSC regulation, and neurodevelopment (**Supplementary Figure 2E and Supplementary Table 1**)^41–43^. Collectively, these findings indicate that our approach detects diverse and disease-relevant SHP2-induced transcription in our model cell line.

### SHP2^WT^ expression shapes the cellular response to EGF stimulation

Next, we examined how cells lacking or expressing SHP2 differ specifically in response to EGF stimulation. SHP2^WT^-transfected cells responded more strongly to EGF stimulation at the transcriptional level than SHP2^KO^ cells (**Figure 2E and Supplementary Table 2**), with changes in gene expression being largely distinct upon EGF-stimulation (**Supplementary Figure 2G**). Gene Set Enrichment Analysis (GSEA) revealed SHP2^WT^-dependent changes in expression of genes involved in proliferative signaling, such as the hallmark mTORc1 and MYC pathways (**Figure 2F**). We also observed an enrichment for genes associated with oxidative phosphorylation and mitochondrial protein degradation pathways (**Figure 2F**).

Interestingly, several early response genes were not significantly differentially expressed as a function of EGF stimulation for SHP2^WT^-expressing cells. Rather, these genes maintain a high basal expression in SHP2^WT^ cells relative to SHP2^KO^ cells, irrespective of stimulation, suggesting that the mere presence of SHP2 produces some basal level of Ras/MAPK signaling (**Supplementary Figure 2H and Supplementary Table 2**). To investigate this further, we determined four unbiased EGF-responsive gene modules based on shared expression patterns across EGF concentrations (**Figure 2G**, **Supplementary Figure 2I**,**J, and Supplementary Table 2**). In SHP2^WT^ cells, at any concentration of EGF, modules 1 and 4 were upregulated and included EGF response genes (1: *ETV4*/7, *CDK4, RHOD* and *PIK3R1;* 4: *EGR1, E2F4, ETF1, POLG*, and *BRD1*) (**Supplementary Table 2**). By contrast, module 2 and 3, which were only downregulated in EGF-stimulated SHP2^WT^ cells, contained several tumor suppressors, such as *NRG1, MAP3K1, CAVIN3, EPHA3/7, CTNNA1*/*3*, and *NOTCH3*. This analysis reveals a broad set of genes co-regulated with SHP2^WT^, but not SHP2^KO^ cells, and highlights the ability of SHP2 to sustain certain EGF signaling markers even without stimulation.

### Pathogenic mutations in SHP2 produce unique gene expression profiles

Having established the transcriptional profile of SHP2^WT^ with and without EGF stimulation, we next examined SHP2 mutant profiles, selecting mutations linked to diverse clinical phenotypes and with varying effects on SHP2 structure (**Figure 3A and Supplementary Table 3**). Two mutations associated with Noonan Syndrome (NS) were included: T42A, which alters N-SH2 binding affinity and specificity^19,44^, and E139D, a C-SH2 mutation that enhances SHP2 basal catalytic activity but does not appear to affect SH2 binding functions^18,19^. Notably, the E139D mutation has also been found in syndromic JMML^20^. E76K, which significantly disrupts auto-inhibition, and T52S, which modestly affects the N-SH2 ligand-binding pocket, were also included as JMML mutations^19,45^. We included NSML mutations Y279C and T468M, which reduce catalytic efficiency while increasing SH2 domain accessibility^5,24,46^. Finally, we included the relatively uncharacterized ALL mutation Q510K, the R138Q mutation found in melanoma and other cancers, which ablates C-SH2 binding capability, and T507K, which disrupts auto-inhibition, alters substrate specificity, and has been observed in neuroblastoma, glioblastoma, and hepatocellular carcinoma^1–3,19,47^.

**Figure 3.**
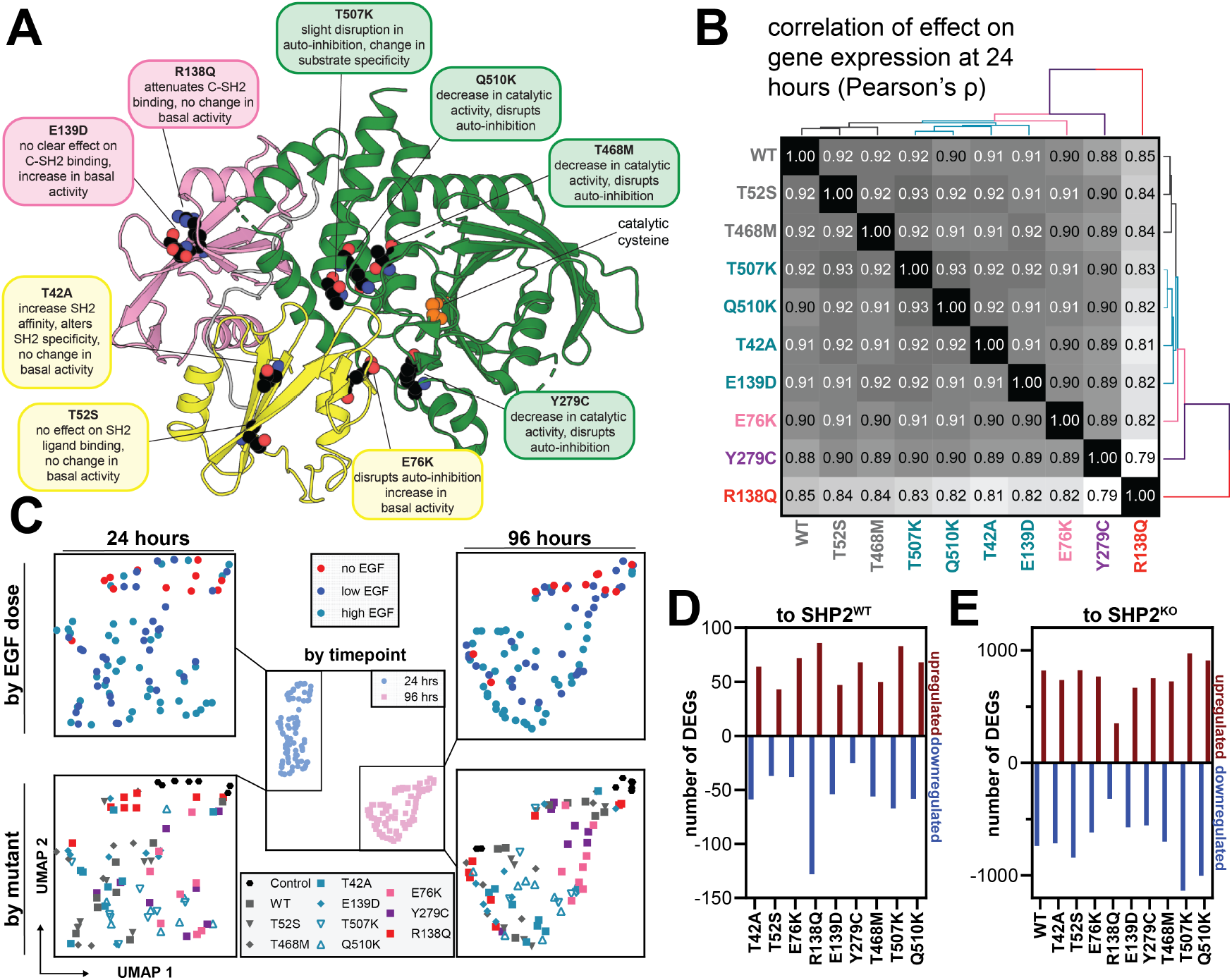
Transcriptomic profiling of SHP2 variants reveals mutational differences. (**A**) Overview of mutants studied in this screen and their position on the protein. Additional descriptions of the mutants are given in **Supplementary Table 3**. (**B**) Heatmap of β coefficient correlation (Pearson’s ρ) with unsupervised hierarchical clustering, comparing SHP2^WT^ and all SHP2 variants at 24 hours. (**C**) Pseudo-bulked log2 fold-change expression of cells grouped by timepoint, SHP2 variant, and EGF dose, against unstimulated SHP2^KO^ cells. Genes were filtered to the union of DEGs across all mutants (5209 genes). Gene space was reduced to 5 principal components, and corrected components were further reduced to 2 Uniform Manifold Approximation and Projection (UMAP)^48^ dimensions for 24 hours (*left*) and 96 hours (*right*). Colors of each mutant represent DEG correlation cluster at 24 hours, as seen in (**B**). (**D**) Number of differentially expressed genes per SHP2 variant, compared to SHP2^KO^, at 24 hours. (**E**) Same as (**D**), but each SHP2 mutant compared to SHP2^WT^.

As with SHP2^WT^, the mutants were expressed in SHP2^KO^ HEK 293 cells, stimulated with a range of EGF concentrations, and harvested at 24 hours and 96 hours (**Figure 2A and Supplementary Figure 1A**). Analysis of differentially expressed genes (DEGs) for each SHP2 variant compared with SHP2^KO^ showed that overall, there is a strong correlation in gene expression changes for all SHP2 variants at respective time points (**Figure 3B, Supplementary Figure 3A, and Supplementary Table 3**). SHP2^R138Q^ was the most distinct mutant at 24 hours, but even this mutant showed a high correlation of effect on gene expression with SHP2^WT^ (Pearson’s ρ of 0.85). At 96 hours both SHP2^R138Q^ and SHP2^Y279C^ were most distinct (Pearson’s ρ of 0.64 and 0.59 with SHP2^WT^, respectively). To visualize the relationship of mutants to each other, we pseudo-bulked (aggregated) the gene expression profiles of cells by time point, SHP2 variant, EGF dose, and replicate, calculated the log_2_ fold-changes to unstimulated SHP2^KO^ cells, and initialized a Uniform Manifold Approximation and Projection (UMAP) embedding with the resulting log_2_ fold-changes (**Figure 3C**). Consistent with the large number of DEGs across mutants upregulated and downregulated (3719 and 3761, respectively, FDR < 0.05) due to stimulation time, time point appears to be the largest determinant of gene expression (**Figure 3C**, *middle panel*). Within each time point, we observed a loose gradient of EGF dose (**Figure 3C**, *top panels*), and separation of SHP2 variants (**Figure 3C**, *bottom panels*, **and Supplementary Figure 3B**).

We next determined a common SHP2-dependent transcriptome, which we defined as genes that are differentially expressed compared to SHP2^KO^ cells (β coefficient < -0.05 or >0.05, false discovery rate < 0.05), shared between at least 5 out of 10 SHP2 variants in our study, and not identified as a DEG for our transfection control (**Supplementary Figure 3C,D**). GSEA of this common transcriptome showed overlap with the previously defined gene sets associated with SHP2^WT^ activity, signifying that different SHP2 mutants drive similar transcriptional programs to SHP2^WT^ and each other (**Supplementary Figure 2A**,**E and Supplementary Tables 1,3**).

Next, we aimed to isolate mutation-dependent changes in transcription. We inspected the top differentially expressed genes between SHP2^WT^ cells and SHP2 mutant cells. We identified between 80 and 214 DEGs across all SHP2 mutants at 24 hours (**Figure 3D**). SHP2^R138Q^ displayed the largest transcriptional differences compared to SHP2^WT^-expressing cells but was transcriptionally more similar to SHP2^KO^ cells compared to all other tested SHP2 variants (**Figure 3E**), suggesting a possible hypomorphic effect at the level of transcription for this SHP2 variant. At 96 hours post-stimulation, the number of DEGs is smaller for both comparison to SHP2^WT^ and to SHP2^KO^ (**Supplementary Figure 3E,F**). Taken together, our initial analysis demonstrates that SHP2 variants are mostly alike, but that differences can be detected.

### EGF-response dynamics are differentially altered by SHP2 mutations

To obtain more specific insights into the mutational differences in SHP2 transcriptomes, we leveraged multi-resolution variational interference (MrVI), a deep generative model that performs sample stratification at single-cell resolution while accounting for technical variability^49^. We recently used MrVI to classify chemical perturbations by their induced transcriptional effects^35^. In this study, we applied the model to detect transcriptional signatures for distinct SHP2 variants (**Figure 4A**). We employed UMAP^48^ for dimension reduction and visualization of cells in the resulting MrVI SHP2 variant/EGF-specific latent space. SHP2^KO^ cells form a distinct cluster separate from all SHP2-containing cells, representing a large driver of variation in our model and further demonstrating the impact that SHP2 presence has on gene expression (**Supplementary Figure 4A**). Thus, to explore more subtle variant specific phenotypes, we continued our analysis in the absence of SHP2^KO^ control cells.

**Figure 4.**
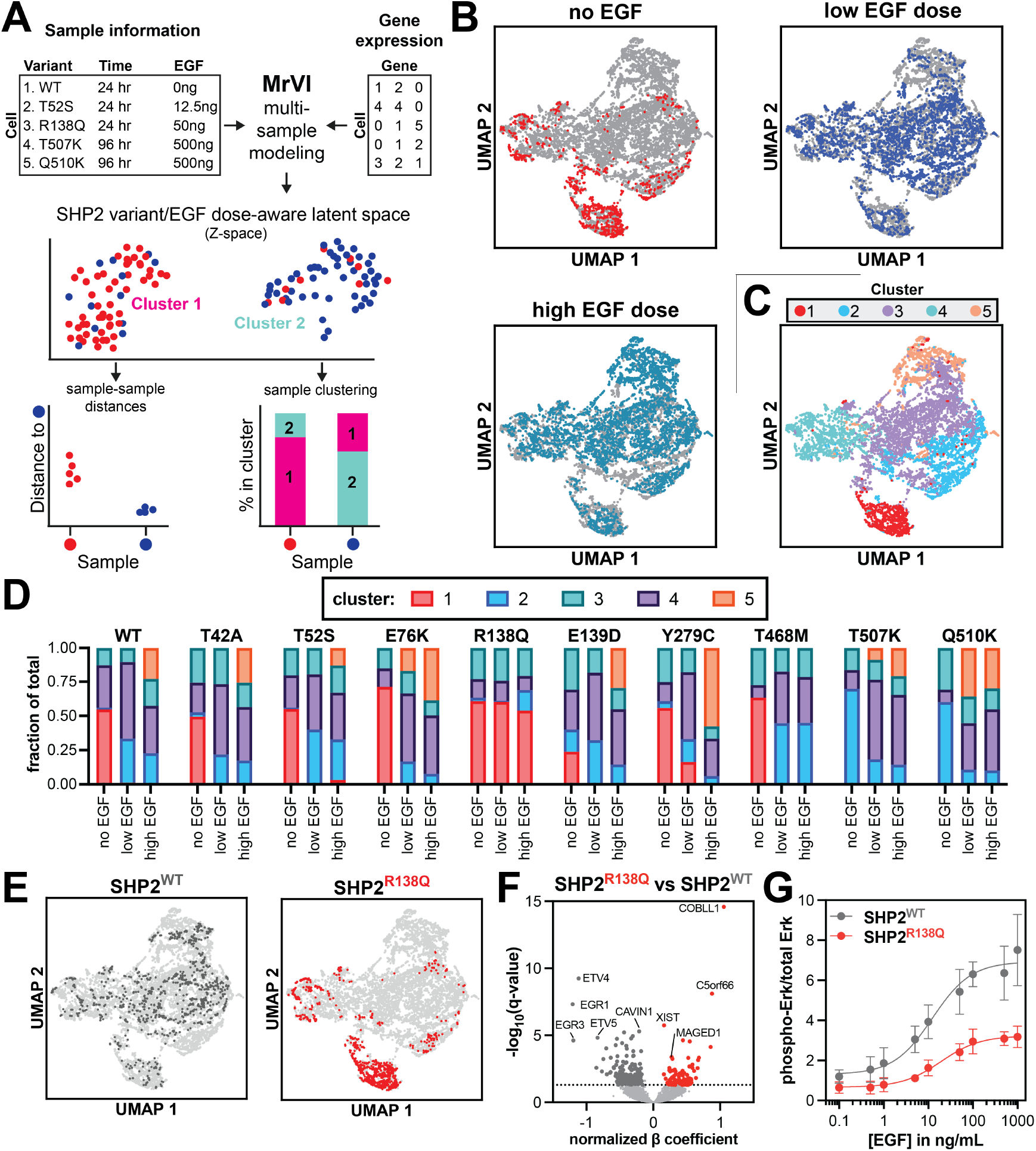
Cellular response to EGF is altered by wild-type and mutant SHP2. (**A**) Overview of MrVI. MrVI model was trained on our 24 hours dataset, in which each combination of SHP2 variant and EGF-dose is defined as the sample-of-origin (96 unique samples), and replicate defined as the technical factor (2 unique replicates). (**B**) UMAP of MrVI Z-space for all single cells (*light grey*), excluding SHP2^KO^ cells. Each respective EGF dose group is indicated per UMAP. (**C**) UMAP of MrVI Z-space for all single cells, excluding SHP2^KO^ cells. Colors indicate clusters as identified by Leiden community detection. (**D**) Bar plots for each SHP2 mutant and their distribution across clusters. (**E**) UMAPs of the MrVI Z-space for SHP2^WT^ and SHP2^R138Q^. (**F**) Volcano plot showing differentially expressed genes between SHP2^R138Q^ and SHP2^WT^ across EGF-concentrations. Significant genes (normalized effect size <-0.15 or >0.15, false discovery rate < 0.05) are labeled in dark grey (SHP2^WT^) and red (SHP2^R138Q^). (**G**) Dose response curves show reduced EGF-response of SHP2^R138Q^ compared to SHP2^WT^. Data points and error bars represent the mean and standard deviation from three independent transfection, stimulation, and blotting experiments.

After omitting SHP2^KO^ cells from our analysis, we noted a separation in the latent space between cells stimulated with no EGF, low concentration of EGF, or high concentrations of EGF (**Figure 4B**). Next, we used Leiden-based community detection^50^ to cluster SHP2-variant expressing cells, resulting in 5 distinct clusters (**Figure 4C and Supplementary Figure 4B**). Analysis of each mutant distribution across these clusters revealed that SHP2^R138Q^ is predominantly present in cluster 1 at any EGF concentration, whereas other SHP2 variants only appear in this cluster in absence of EGF stimulation (**Figure 4D,E and Supplementary Figure 4C,D**). Furthermore, SHP2^R138Q^ and SHP2^T468M^ never occupy cluster 5, and SHP2^WT^ and several SHP2 variants (SHP2^T42A^, SHP2^T52S^, SHP2^E139D^, SHP2^Y279C^) only appear in cluster 5 when stimulated with high doses of EGF (**Figure 4D and Supplementary Figure 4D**). By contrast, SHP2^E76K^, SHP2^T507K^ and SHP2^Q510K^ already populate cluster 5 at low EGF doses. Similar trends were observed when analyzing the counterfactual cell distances determined by MrVI, comparing each mutant and stimulation condition to unstimulated SHP2^WT^ cells (**Supplementary Figure 4E**). These observations demonstrate how different structural perturbations to SHP2 can alter its EGF-responsiveness.

One notable conclusion from our comparison of SHP2 mutants is that SHP2^R138Q^-expressing cells at any dose of EGF behave most similarly to unstimulated cells expressing almost any other SHP2 variant. Analysis of the differentially expressed genes between SHP2^WT^ and SHP2^R138Q^ revealed that SHP2^R138Q^ cells do not express canonical EGF-response genes, such as *EGR1*/*3*, to the same extent as SHP2^WT^ cells (**Figure 4F**). We previously showed that SHP2^R138Q^ has an almost non-functional C-SH2 domain^19^, and co-localized proteins are less likely to be tyrosine-phosphorylated when compared with SHP2^WT^-colocalized proteins^26^. C-SH2/phosphoprotein interactions play an important role in localizing SHP2 to signaling complexes^51^, and SHP2^R138Q^ may thus be unable to interact with EGFR pathway phosphoproteins, thereby decreasing responsiveness to EGF stimulation. Consistent with this, EGF-induced changes in gene expression with SHP2^R138Q^ are much smaller than with SHP2^WT^ and do not include canonical EGF response genes (**Supplementary Figure 4F**). Furthermore, when we examined Erk phosphorylation as a marker of EGF signaling, we observed a reduced EGF-dependent phospho-Erk levels in SHP2^R138Q^-expressing cells when compared to SHP2^WT^-expressing cells, with a modest shift in EC_50_ for EGF and large reduction in signal amplitude (**Figure 4G and Supplementary Figure 4G**).

Some of the proliferative, EGF-response genes that were depleted with SHP2^R138Q^ expression, such as *EGR1* and *ETV4*, are also known cancer-associated genes^52^. To understand which genes might drive the oncogenicity associated with the R138Q mutation, we compared our gene expression profile to genes known to be associated with cancers where SHP2^R138Q^ has been observed, including melanoma and prostatic adenocarcinoma^52^. *XIST*, a known regulator of malignant melanoma, was significantly enriched in the SHP2^R138Q^ transcriptome^53^, as was C5orf66, a long non-coding RNA, which can function as both an oncogene and tumor suppressor dependent on tissue type^54–57^. We also identified *MAGED1* (melanoma-associated antigen 1), a member of the MAGE family which is frequently upregulated in melanoma and other cancers and is a therapeutic target^58^. However, over-expression of *MAGED1* can suppress cell cycle progression and tissue invasion in other cell systems^59^. The most upregulated gene in SHP2^R138Q^-expressing cells, considering both SHP2-driven effects and EGF-induced effects, was *COBLL1* (**Figure 4F and Supplementary Figure 4F**), which is involved in the oncogenesis of prostate cancer and chronic lymphocytic leukemia^60,61^. Thus, while the SHP2^R138Q^ mutant has a severely attenuated response to EGFR activation, its expression can still upregulate known oncogenes.

### SHP2^T507K^ and SHP2^Q510K^ drive Ras/MAPK signaling in unstimulated cells

Whereas SHP2^R138Q^ was unique in the extent to which it attenuates EGF responsiveness (**Figure 4E,F**), two mutations on the catalytic Q-loop (**Supplementary Figure 5A**), SHP2^T507K^ and SHP2^Q510K^, were unique from all other SHP2 variants in that they did not occupy cluster 1 in the absence of EGF stimulation (**Figure 4D and Figure 5A**). Instead, these variants appear to drive an altered basal cellular state that is most represented by cluster 2 (**Figure 4E**). The T507K mutation, which has been biochemically characterized^47^, is associated with several solid tumors, including hepatocellular carcinoma, glioblastoma, and neuroblastoma^1–3^. By contrast, the relatively unstudied Q510K is mainly associated with ALL, although it has also been observed in solid tumours^62^.

**Figure 5.**
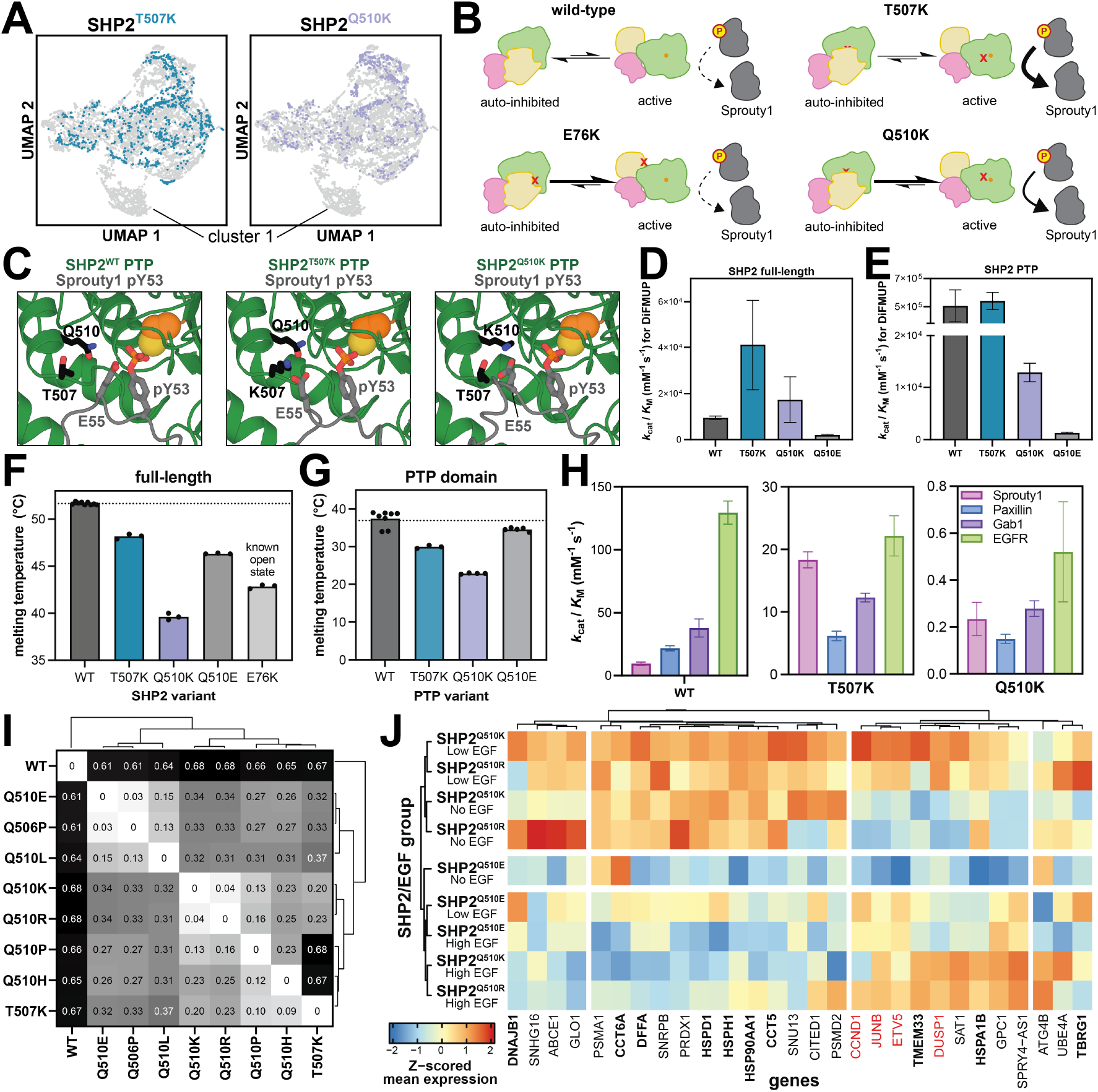
Q-loop mutations can alter substrate specificity and conformational stability to modulate downstream transcription. (**A**) UMAP of the MrVI Z-space for SHP2^T507K^ and SHP2^Q510K^ shows absence of cells in cluster 1. (**B**) Schematic showing the structure and activity changes in SHP2^Q510K^ relative to SHP2^WT^, SHP2^E76K^, and SHP2^T507K^. The Q510K mutation shifts the protein towards the open conformation, while also enhancing Sprouty1 dephosphorylation. (**C**) AlphaFold 3 models of SHP2^T507K^ (top) and SHP2^Q510K^ (bottom), bound to Y53-phosphorylated Sprouty1 in the active site, showing the proximity of K507 and K510 with E55 on Sprouty1. (**D**) Catalytic efficiencies of full-length SHP2^WT^, SHP2^T507K^, and SHP2^Q510K^ against DiFMUP. (**E**) Same as (**D**), but for the isolated PTP domains. (**F**) Melting temperatures for full-length SHP2^WT^, SHP2^T507K^, SHP2^Q510K^, and SHP2^Q510E^. SHP2^E76K^, a known open conformation mutant, is shown for reference. (**G**) Same as (**F**), but for isolated PTP domains. (**H**) Dephosphorylation assay with PTP^WT^, PTP^T507K^, and PTP^Q510K^ showing switch in substrate preferences. Full peptide sequences are indicated in the Methods. (**I)** Pairwise MrVI counterfactual cell distances, showing the largest distances between SHP2^WT^ and any Q-loop mutant. SHP2^Q510K^ and SHP2^Q510R^ show the smallest distance observed. (**J**) Heatmap for differentially expressed genes for SHP2^Q510K/R^ versus SHP2^WT^. Z-scored mean expression for SHP2^Q510K^, SHP2^Q510R^, and SHP2^Q510E^ are visualized. Gene names in bold represent chaperone and protein folding genes. Red gene names represent EGF response genes.

We hypothesized that these mutations converge in their gene expression profiles due to a shared molecular mechanism (**Figure 5B**). Specifically, SHP2^T507K^ is known to have reduced catalytic activity against many phosphopeptide substrates, however, due to the introduction of a positive charge in the substrate-binding pocket, SHP2^T507K^ has higher than wild-type level catalytic efficiency for substrates with a complimentary acidic residue, such as Sprouty1 pY53 ^47^. This change in substrate-preferences has been linked to T507K-pathogenic signaling, as Sprouty1 is a negative regulator of Ras, and its dephosphorylation by SHP2 causes activation of the Ras/MAPK pathway^47^. Furthermore, T507K modestly destabilizes the auto-inhibited state of SHP2, enhancing its propensity for activation by phosphoprotein binding^47^. Q510 is a key catalytic residue, and mutations at this site, including Q510K, impair catalysis^17,21^. However, structural models suggest that SHP2^T507K^ and SHP2^Q510K^ might have similarly remodeled active site electrostatics, which could result in similar changes in substrate-specificity (**Figure 5C**)^63^.

To test if T507K and Q510K dysregulate SHP2 through similar molecular mechanisms, we measured the catalytic activities of full-length and isolated phosphatase (PTP) domain constructs of SHP2^WT^, SHP2^T507K^, and SHP2^Q510K^. Full-length SHP2^WT^ and SHP2^Q510K^ have comparable activity against the fluorogenic model substrate DiFMUP, while SHP2^T507K^ shows a large increase in catalytic efficiency, consistent with a previous study on SHP2^T507K^ (**Figure 5D**)^47^. For the isolated PTP domains, we found that PTP^Q510K^ was substantially less active than PTP^WT^ or PTP^T507K^ against DiFMUP (**Figure 5E**). This discrepancy between full-length and wild-type proteins could be explained by the ability of the Q510K mutation to disrupt auto-inhibition, thereby compensating for the loss of a catalytic residue (**Figure 5B**). Indeed, differential scanning fluorimetry demonstrated that SHP2^Q510K^ had a dramatically lower melting temperature than SHP2^WT^ and SHP2^T507K^, indicative of a more open conformation (**Figure 5F**)^17,19,64^. Even in the context of the isolated PTP domain, the Q510K mutation showed a much lower melting temperature than PTP^WT^ (**Figure 5G**), suggesting that this mutant not only disrupts auto-inhibition in the full-length protein but also intrinsically destabilizes the isolated PTP domain.

Next, we measured the activity of the isolated PTP domains against 4 peptide-substrates: Paxillin pY118, Sprouty1 pY53, Gab1 pY589, and EGFR pY992, which were previously used to profile the change in SHP2^T507K^ substrate preferences^47^. Consistent with previously reported results, we saw an increase in preference for Sprouty1 for PTP^T507K^ when compared to PTP^WT^ (**Figure 5H and Supplementary Figure 5B**). Notably, while PTP^Q510K^ overall shows strong catalytic impairment, we observe an increased preference for Sprouty1, suggesting that the Lys-substitutions on the two nearby sites have convergent effects on substrate preferences (**Figure 5H and Supplementary Figure 5B**). Furthermore, the destabilizing effect of the Q510K mutation disrupts auto-inhibition to such an extent that the full-length protein has comparable activity to SHP2^WT^ (**Figure 5B,D**). While substrates other than Sprouty1 may be at play for either mutant, and other structural explanations might also be relevant, our biochemical and transcriptomic results suggest that the Q510K and T507K operate at least partly through similar mechanisms.

### The identity of the Q510 substitution fine-tunes functional outcomes

In addition to T507K and Q510K, there are several other pathogenic mutations in the Q-loop of SHP2 (**Supplementary Figure 5A**). Particularly, Q510 has several other known pathogenic substitutions (Q510H/E/L/P/R) – all of which are one nucleotide away from the wild-type sequence and have distinct disease outcomes (**Supplementary Table 3**). With the exception of Q510K, all observed Q510 mutations are associated with NSML, with Q510L/P/R also having been implicated in NS. Like Q510K, Q510E is also associated with ALL, whereas Q510P, L, and H have been found in AML patients. Additionally, all Q510 mutants occur in solid cancers. While loss of the wild-type residue may fully explain dysregulatory effects of mutations at a particular site, evidence from other proteins such as Ras GTPases suggests that the identity of the substituted amino acid can also dictate functional outcomes^65,66^. Indeed, for SHP2, we see differences in basal activity and stability for Q510K and Q510E that could have downstream consequences (**Figure 5D-G**). Thus, we conducted another transcriptomic screen focused on Q-loop mutations, including all disease-relevant Q510 substitutions, T507K, and another common mutation at a catalytic residue, Q506P (**Supplementary Figure 6A-D**). All Q-loop mutants appeared distinct from SHP2^WT^ and induced a similar number of differentially expressed genes relative to SHP2^WT^ (**Supplementary Figure 6E**,**F**).

Building on the observation from our previous screen that SHP2^T507K^ and SHP2^Q510K^ were most distinct from other SHP2 variants in unstimulated conditions, we trained a specific MrVI model on the unstimulated cells at 24 hours. Then, we calculated the counterfactual cell distances between different SHP2 variants in this screen. Consistent with the trend found in our correlation of β coefficients (**Supplementary Figure 6E**), we found that SHP2^WT^ shows the largest distance to any of the mutants in our screen (distance = 0.61 – 0.68) (**Figure 5I**). Furthermore, we found that SHP2^Q510K^ and SHP2^Q510R^, which introduce a positive charge, are remarkably similar (distance = 0.04), whereas SHP2^Q510E^, which brings about a negative charge, is the most distant mutant from SHP2^Q510K/R^ (distance = 0.34) (**Figure 5I**).

To understand what drives these trends, we analyzed differentially expressed genes between SHP2^WT^ and the group of SHP2^Q510K/R^. We identified several EGF response genes, such as *JUNB, CCND1, DUSP1* and *ETV5*, as increasingly expressed in cells with the SHP2^Q510K/R^, many of which were not upregulated to the same extent with SHP2^Q510E^ (**Figure 5J**, *red genes*, **and Supplementary Table 3**). This suggests that SHP2^Q510R^ potentially shares the altered cell state that we previously observed for SHP2^Q510K^ and SHP2^T507K^, which could be explained by a similar charge-based change in substrate specificity. By contrast, SHP2^Q510E^ is effectively catalytically dead but has a mildly destabilized auto-inhibited state, somewhere between SHP2^WT^ and SHP2^Q510K^ (**Figure 5D-G**). Thus, it can only drive signaling through its scaffolding functions. Notably, SHP2^Q510E^ is most similar to SHP2^Q506P^ in our transcriptomics data (**Figure 5I**). This mutation also reduces catalytic activity, and it has been reported to destabilize SHP2 auto-inhibition to the same extent as SHP2^Q510E 21^.

Finally, we were surprised to find that the intersection of the SHP2^Q510K^ and SHP2^Q510R^ data showed enrichment for genes encoding chaperones and other proteostasis machinery (**Supplementary Table 3**). We compared the Z-scored mean expression of these identified genes in SHP2^Q510K/R^ data to SHP2^Q510E^ data and found that these genes are expressed to much lesser degree in SHP2^Q510E^ (**Figure 5J**, *bolded genes*). One plausible explanation for this difference is that the Q510R mutation, similar to Q510K, may destabilize the PTP domain of SHP2, triggering proteostasis machinery. By contrast, Q510E is much less destabilizing, both for full-length SHP2 and the isolated PTP domain (**Figure 5F,G**), which may explain why a similar response is not observed in SHP2^Q510E^-expressing cells. Overall, our data suggest that the distinct substitutions at Q510 can have diverse effects on protein conformation, stability, and activity, which are likely to shape unique downstream signaling and transcriptional programs.

## Discussion

In this project, we conducted two multiplexed single-cell transcriptomic screens with cells expressing a variety of pathogenic SHP2 mutants, stimulated with a range of EGF doses across multiple time points. First, by comparing SHP2^WT^-expressing cells directly to SHP2^KO^ cells, we show that the presence of SHP2 is essential for expression of EGF-response genes, such as *EGR1/3* and *ETV4/5*. As a result, SHP2^KO^ cells are defective in manifesting a cellular response to EGF stimulation. SHP2^WT^-expressing cells showed upregulation of several key cell signaling pathways, including mTORC1, and MYC pathways. Interestingly, the SHP2^KO^ cells only showed upregulation of FGFR3-related signaling, which is consistent with previous studies showing that SHP2 inhibition led to compensatory activation of FGFR signaling and rebound ERK activity, suggesting that in cells navigate the absence of SHP2 by escaping to other cell signaling pathways^67,68^.

Next, we compared SHP2^WT^ to nine SHP2 mutants, chosen for their range of effects on protein structure and activity, along with diverse disease contexts. We observed a distinctive correlation between the protein-level mechanism of dysregulation and resulting cell states. For example, SHP2^R138Q^, which has a defective C-SH2 domain^19^, attenuated proximal signaling in response to EGF (lower Erk phosphorylation) and led to significantly diminished transcription of EGF-response genes. Importantly, SHP2^R138Q^ retains normal catalytic activity, both in basal conditions as well as when the N-SH2 domain is engaged by a phosphopeptide^19^, illustrating how non-catalytic properties of SHP2 are critical for EGF signaling. Previous work has shown that catalytically-dead SHP2^C459S^ is unable to activate the Ras/MAPK pathway in response to EGF stimulation^69^. This suggests that both the C-SH2 binding function and phosphatase domain catalytic activity of SHP2, are needed for activation of the Ras/MAPK pathway in response to EGF stimulation, highlighting the importance of the scaffolding function of SHP2, as well as its interplay with catalytic activity. Interestingly, our transcriptomics data show that the gene expression profile mediated by SHP2^R138Q^ is distinct from SHP2^KO^, indicating that some of its signaling functions are intact, and novel oncogenes are overexpressed that are not seen in the SHP2^KO^ or SHP2^WT^ context.

Our transcriptomics data also revealed surprising instances of functional convergence by two apparently unrelated mutations and functional divergence by different substitutions at the same site. Specifically, we found that the unstudied cancer mutant SHP2^Q510K^ partly phenocopies SHP2^T507K^ by introducing a lysine into the substrate-binding pocket of the phosphatase domain and altering substrate specificity. As a result, both of these mutants mediate similar EGF-dependent transcriptional responses that are distinct from all other mutants in our screen. By contrast, other pathogenic mutations at Q510 unexpectedly showed a range of downstream effects. SHP2^Q510K^ and SHP2^Q510R^, which both introduce a positive charge in the active site, produce the most similar gene expression profiles, whereas we found that SHP2^Q510E^, which introduces a negative charge, has distinct effects on protein stability, auto-inhibition, and catalytic activity, resulting in a divergent transcriptome. For many known missense mutations in SHP2, it appears that the loss of the original residue is more detrimental to protein function than the identity of the new amino acid. For example, while SHP2^E76K^ is well-known and well-studied, substitutions to D, G, A, Q, V, and M at this position have also been identified in patients^70^, and all of these substitutions disrupt autoinhibition to hyperactivate SHP2^17,20,62^. By contrast, for Q510 we find that the identity of the resulting mutation at a site can also dictate downstream functions. This “original-centric” view of pathogenic mutations is being challenged in other systems as well, including the well-studied oncogene Ras, where different G12 mutations have distinct effects on protein interactions and GTP hydrolysis rates^17,20,62^.

A critical feature of our experimental design that yielded the aforementioned insights is that we conducted these screens in a homogenous genetic background, SHP2^KO^ HEK 293 cells. This approach isolates how structural and biochemical consequences of mutations in SHP2 propagate to changes in cell state, without other confounding factors, including genetic, transcriptomic, or proteomic variation. Through this approach, we were able to amplify mutant-specific outcomes and connect changes in cell state to nuanced mutant-dependent changes in SHP2 structure, stability, and molecular recognition. We acknowledge that the mutants in our study occur in a broad range of human diseases and thus affect a broad range of cell types. In the congenital disorders Noonan Syndrome and Noonan Syndrome with Multiple Lentigines, SHP2 mutations are inherited and systemic, whereas somatic SHP2 mutations in cancers are localized to specific cell and tissue types. Thus, these mutations naturally drive diseases in a wide array of cellular and mutational contexts. Our reductionist approach provides a baseline for connecting SHP2 structural perturbations to cellular outcomes and lays the foundation for deeper mechanistic studies in disease-relevant cell lines, animal models, or patient samples.

In our study, we focused on just over a dozen pathogenic mutations that have varied effects on SHP2 at the molecular level. By analyzing how these diverse mutants influence the transcriptome, we were able to show that corresponding changes to the conformational state of SHP2, to non-catalytic protein-protein interactions, and even to its substrate specificity, can propagate into major differences in cellular outcomes, separate from effects coming from basal catalytic activity. Our results highlight the bidirectional value of integrating structural and cellular data: consideration of protein structure and biochemistry can inspire insightful cellular experiments, while unbiased assessments of cellular phenotypic effects—such as through transcriptomics—can, in turn, reveal unexpected biochemical insights. In the future, one can envision taking a more expansive and unbiased approach to gain even deeper insights. The analysis of comprehensive scanning mutagenesis libraries, coupled with functional selection and deep sequencing, is yielding new insights into protein stability, regulation, molecular recognition, catalysis, and drug resistance^71^. Combining these deep mutational scanning approaches with multiplexed single-cell transcriptomics could yield a powerful framework for mapping the effects of mutations from the molecular to the cellular scale.

## Acknowledgements

We would like to thank the members of the Shah and McFaline-Figueroa labs for their scientific insights and helpful discussions, in particular Nicholas Hou for MrVI guidance. This research was funded by NIH/NIGMS grant R35GM138014 to NHS. J.L.M.-F acknowledges support from grants from the NIH (R35HG011941) and the NSF (2146007). These studies used the resources of the Cancer Center Sequencing Core Facility at Columbia University funded in part through Center Grant P30CA013696.

## Author Contributions

AEV conceived, designed, performed, analyzed, and interpreted the experiments; performed statistical analysis; and wrote the manuscript. RMG designed, performed, analyzed and interpreted the experiments, performed statistical analysis and edited the manuscript. ZJ performed, analyzed and interpreted experiments. ML synthesized key reagents. JLMF designed, analyzed and interpreted the experiments and edited the manuscript. NHS designed, analyzed, and interpreted the experiments, and wrote the manuscript.

## Competing interests

The authors declare no competing interests.

## Data and materials availability

Raw and processed data can be accessed and downloaded from NCBI GEO under accession number Series GSE300865. The code necessary to reproduce the analyses in this study can be found at Github https://github.com/mcfaline-figueroa-lab/sci-Plex-SHP2.

## Supplementary Information

### Supplementary Information file contains

Supp. Fig. 1. Comprehensive screen of pathogenic PTPN11 mutations using sci-Plex

Supp. Fig. 2. Comparison of SHP2^WT^ transcriptome to SHP2^KO^

Supp. Fig. 3. Analysis of shared and distinct SHP2 variant transcriptomes

Supp. Fig. 4. MrVI analysis of SHP2 variant response to EGF stimulation

Supp. Fig. 5. Michaelis-Menten analysis of PTP^T507K^ and PTP^Q510K^

Supp. Fig. 6. Screening of Q-loop mutants Materials and Methods

Supplementary tables can be found as separate spreadsheet files:

Supp. Table 1. SHP2-driven effects in gene expression

Supp. Table 2. EGF-driven effects in gene expression

Supp. Table 3. Gene expression of SHP2 mutants

## Notes

### Competing Interest Statement

The authors have declared no competing interest.

