## Supplementary Information for "Multiplexed single-cell transcriptomics reveals diverse phenotypic outcomes for pathogenic SHP2 variants"

#### Table of contents

| | | |
|---|---|---|
| Supp. Fig. 1 | Comprehensive screen of pathogenic *PTPN11* mutations using sci-Plex | pg 2 |
| Supp. Fig. 2. | Comparison of $SHP2^{WT}$ transcriptome to $SHP2^{KO}$ | pg 3 |
| Supp. Fig. 3. | Analysis of shared and distinct SHP2 variant transcriptomes | pg 4 |
| Supp. Fig. 4. | MrVI analysis of SHP2 variant response to EGF stimulation | pg 5 |
| Supp. Fig. 5. | Michaelis-Menten analysis of $PTP^{T507K}$ and $PTP^{Q510K}$ | pg 6 |
| Supp. Fig. 6. | Screening of Q-loop mutants | pg 7 |
| Materials and methods | | pg 8 |
| Supplementary references | | pg 13 |

**Supplementary Tables** (included as separate spreadsheet files)

Table 1: SHP2-driven effects in gene expression
- a) Differentially expressed genes test (SHP2 coefficients) for $SHP2^{WT}$ vs $SHP2^{KO}$;
- b) Gene Ontology analysis of DEGs upregulated in $SHP2^{WT}$;
- c) Gene Ontology analysis of DEGs downregulated in $SHP2^{WT}$;
- d) GSEA analysis of $SHP2^{WT}$ vs $SHP2^{KO}$.

Table 2: EGF-driven effects in gene expression
- a) EGF-induced differentially expressed genes for each sample;
- b) Gene Modules.

Table 3: Gene expression of SHP2 mutants
- a) Structural, biochemical, and clinical effects ofSHP2 variants;
- b) Differentially expressed gene test for all SHP2 variants vs $SHP2^{KO}$;
- c) Differentially expressed gene test for all SHP2 mutants vs $SHP2^{WT}$;
- d) Differentially expressed genes test for $SHP2^{Q510K/R}$ vs $SHP2^{WT}$.

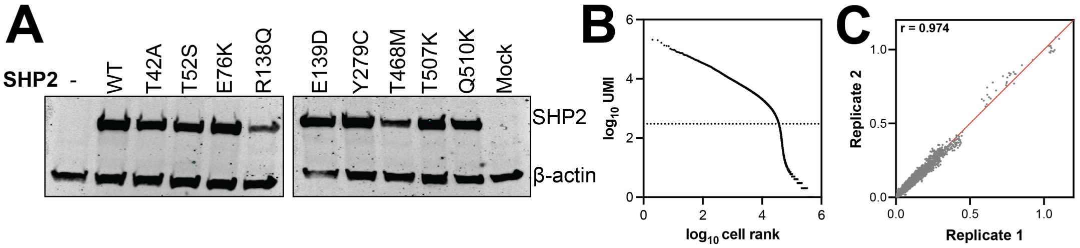


D

| Timepoint | Total Cell Count | Mean Cell Count per SHP2mut-EGF | Max Cell Count per SHP2mut-EGF | Max Cell Count per SHP2mut-EGF | Median UMI | Median Hash UMI |
|---|---|---|---|---|---|---|
| 24hr | 13,446 | 140.1 | 265 | 64 | 926 | 48 |
| 96hr | 16,270 | 169.5 | 413 | 22 | 878 | 58 |
| Total | 29,716 | 154.8 | 413 | 22 | 900 | 53 |

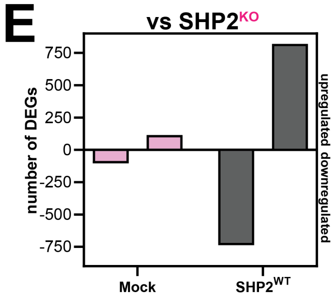


**Supplementary Figure 1. Comprehensive screen of pathogenic *PTPN11* mutations using sci-Plex.** (**A**) Western blot showing expression of $SHP2^{WT}$ and mutants transfected into $SHP2^{KO}$ HEK293 cells. (**B**) Knee plot of unique molecular identifier (UMI) vs cell rank with a UMI cut-off of 300 to filter for high-quality cells (**C**) Correlation of the mean expression between two independent replicates at 24 hour (*left*) and 96 hours (*right*). (**D**) Table of experimental summary metrics. (**E**) Number of differentially expressed genes (normalized effect size <-0.25 or >0.25, false discovery rate <0.05) for mock-transfected cells and $SHP2^{WT}$-expressing cells respectively, compared to $SHP2^{KO}$ cells. Number of differentially expressed genes for SHP2 variants vs $SHP2^{KO}$ can be found in **Figure 3E**.

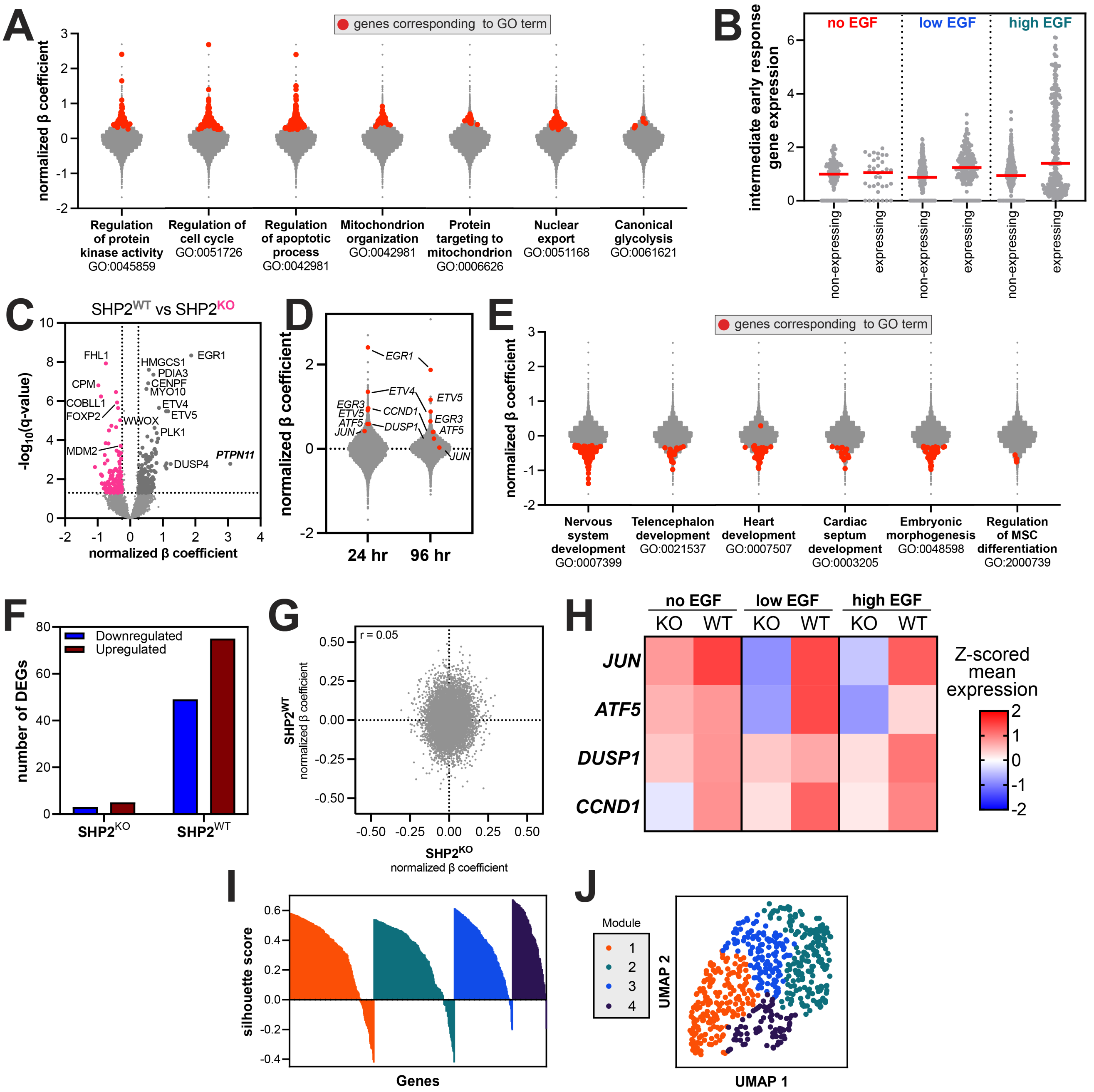


**Supplementary Figure 2. Comparison of SHP2$^{WT}$ transcriptome to SHP2$^{KO}$.** (**A**) Normalized effect sizes for differentially expressed genes (SHP2$^{WT}$ vs SHP2$^{KO}$; false discovery rate < 0.05, normalized effect size > 0.25; red dots) corresponding to statistically enriched Gene Ontology (GO) terms. Genes related to transcription, translation, and RNA more broadly were removed prior to analysis. (**B**) Expression of immediate early response genes of non-SHP2 expressing cells in the SHP2$^{WT}$ sample versus SHP2$^{WT}$-expressing cells. (**C**) Volcano plots showing SHP2-induced differentially expressed genes for SHP2$^{KO}$ and SHP2$^{WT}$ at 96 hours. Any significant transcript with a normalized effect size of > 0.25 or <-0.25 is colored. (**D**) Comparison of early response genes (*EGR1/3, ETV4/5, DUSP1, CCND1, ATF5, JUN*; red dots) at 24 hours and 96 hours between SHP2$^{WT}$ (top of violin) and SHP2$^{KO}$ (bottom of violin). (**E**) Same as (**A**), but for differentially expressed genes with false discovery rate < 0.05, normalized effect size < -0.25. MSC = mesenchymal stem cells. (**F**) Number of up- and down-regulated genes for SHP2$^{KO}$ and SHP2$^{WT}$. The number of DEGs is larger for SHP2$^{WT}$. (**G**) Correlation of β coefficients for differentially expressed genes between SHP2$^{KO}$ and SHP2$^{WT}$. Low β coefficient shows SHP2$^{KO}$ and SHP2$^{WT}$ are not correlated or inversely correlated but rather have their own gene expression effects. (**H**) Heatmap showing expression of early response genes with high basal expression in SHP2$^{WT}$ unstimulated cells compared with SHP2$^{KO}$ cells. (**I**) Silhouette plot of gene modules suggests that 4 models achieves good consistency within clusters and separation between clusters for most genes. (**J**) UMAP of gene space with 4 gene modules indicated by color.

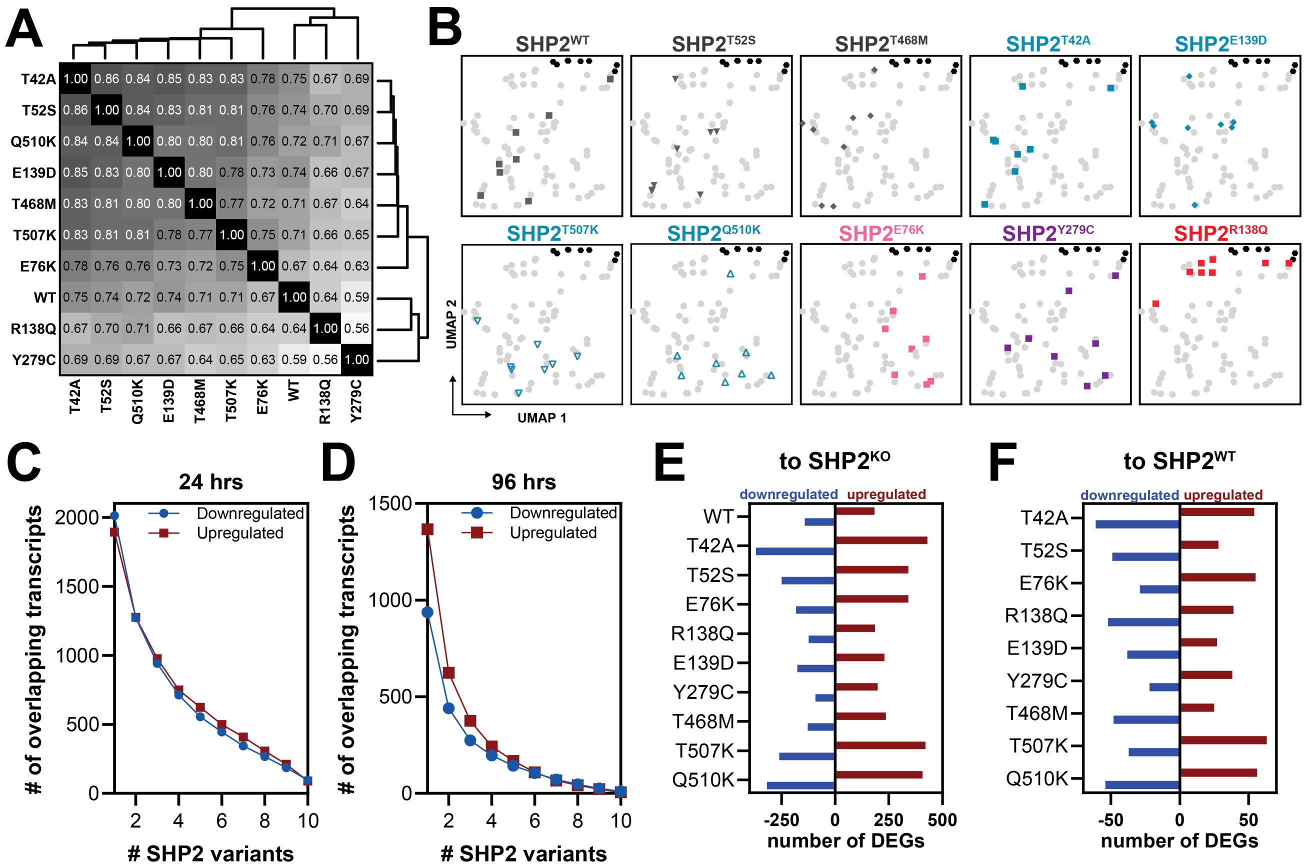


**Supplementary Figure 3. Analysis of shared and distinct SHP2 variant transcriptomes.** (**A**) Heatmap of β coefficient correlation (pearson's r) with unsupervised hierarchical clustering, comparing SHP2$^{WT}$ and all SHP2 variants at 96 hours. SHP2$^{Y279C}$-expressing cells appear most distinct. (**B**) Pseudo-bulked $\log_2$ fold-change expression of cells grouped by timepoint, SHP2 variant and EGF dose against unstimulated SHP2$^{KO}$ cells. (**C**) Number of overlapping genes that are significantly enriched (normalized effect size <-0.25 or >0.25, and false discovery rate < 0.05) over the SHP2$^{KO}$ control at 24 hours. (**D**) Same as (**C**), but for 96 hours. (**E**) Number of differentially expressed genes (normalized effect size <-0.25 or >0.25, and false discovery rate < 0.05) per SHP2 mutant, compared to SHP2$^{KO}$, at 96 hours after EGF stimulation. (**F**) Same as (**E**), but for each SHP2 variant against SHP2$^{WT}$.

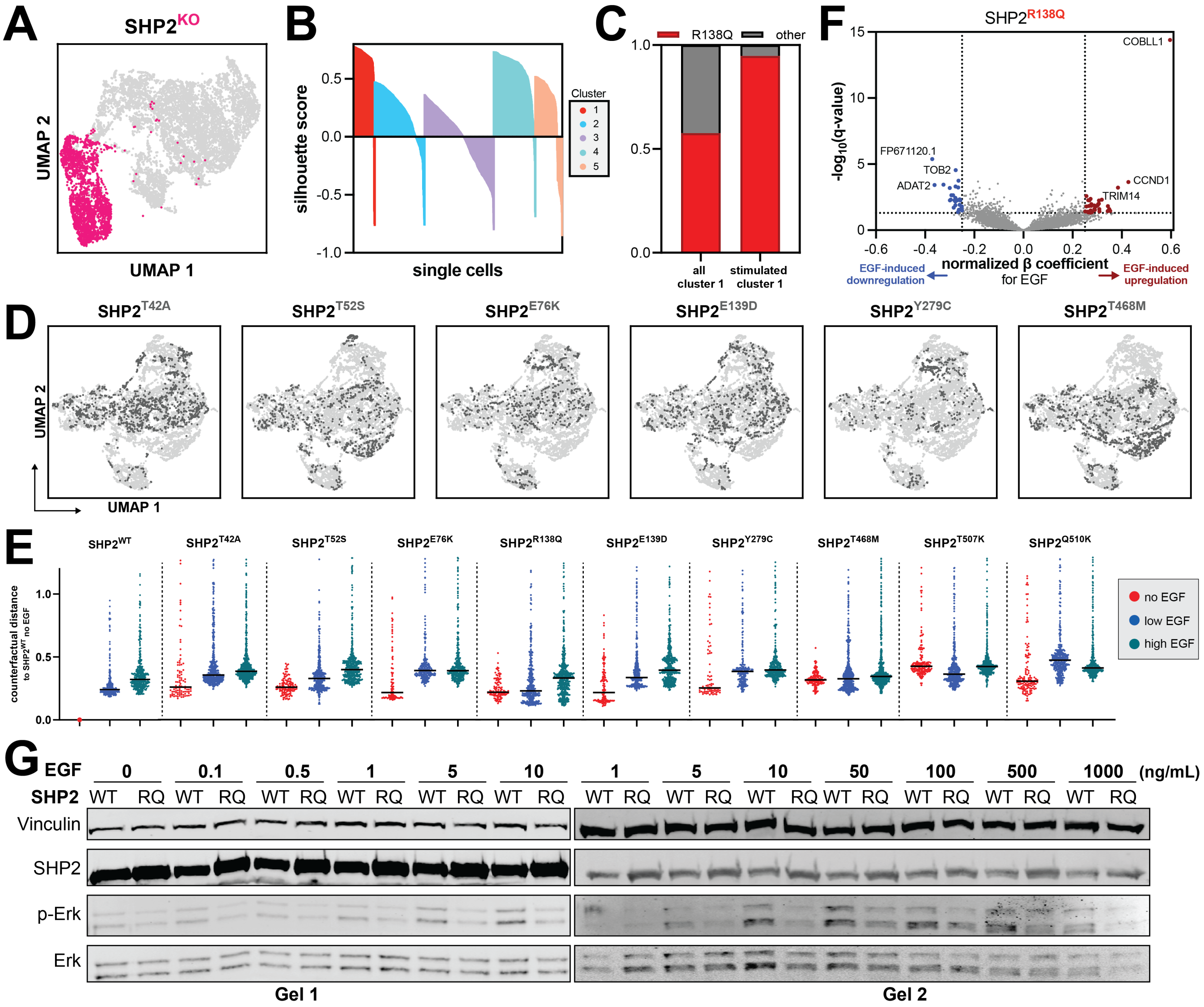


**Supplementary Figure 4**. **MrVI analysis of SHP2 variant response to EGF stimulation.** (**A**) UMAP of the MrVI Z-space for single cells shows distinct grouping for SHP2$^{KO}$ (*pink*) compared to any SHP2 variant (*light grey*). (**B**) Silhouette plot of Leiden-based clustering in cell space suggests that 5 clusters achieves good consistency within clusters and separation between clusters for most cells. (**C**) Mutant per cluster distribution for cluster 5 demonstrates disproportionate representation of SHP2$^{R138Q}$. (**D**) UMAPs of MrVI Z-space for cells expressing different SHP2 variants (*dark grey*). (**E**) Distribution of MrVI U-space to Z-Space distance between each mutant and unstimulated SHP2$^{WT}$ cells shows distinct behaviors in response to EGF dose: positively correlated (SHP2$^{WT}$, SHP2$^{T42A}$, SHP2$^{T52S}$, SHP2$^{E139D}$), early saturating (SHP2$^{E76K}$, SHP2$^{Y279C}$), desensitized (SHP2$^{R138Q}$), unaffected (SHP2$^{T468M}$), and irregular (SHP2$^{T507K}$, SHP2$^{Q510K}$). (**F**) Volcano plot showing EGF-induced differentially expressed genes for SHP2$^{R138Q}$. Any significant (false discovery rate < 0.05) transcript with a normalized effect size (NES) of > 0.25 or <-0.25 is colored. (**G**) Representative western blot of dose-response curves for SHP2$^{WT}$ and SHP2$^{R138Q}$ (n = 3 independent transfections).

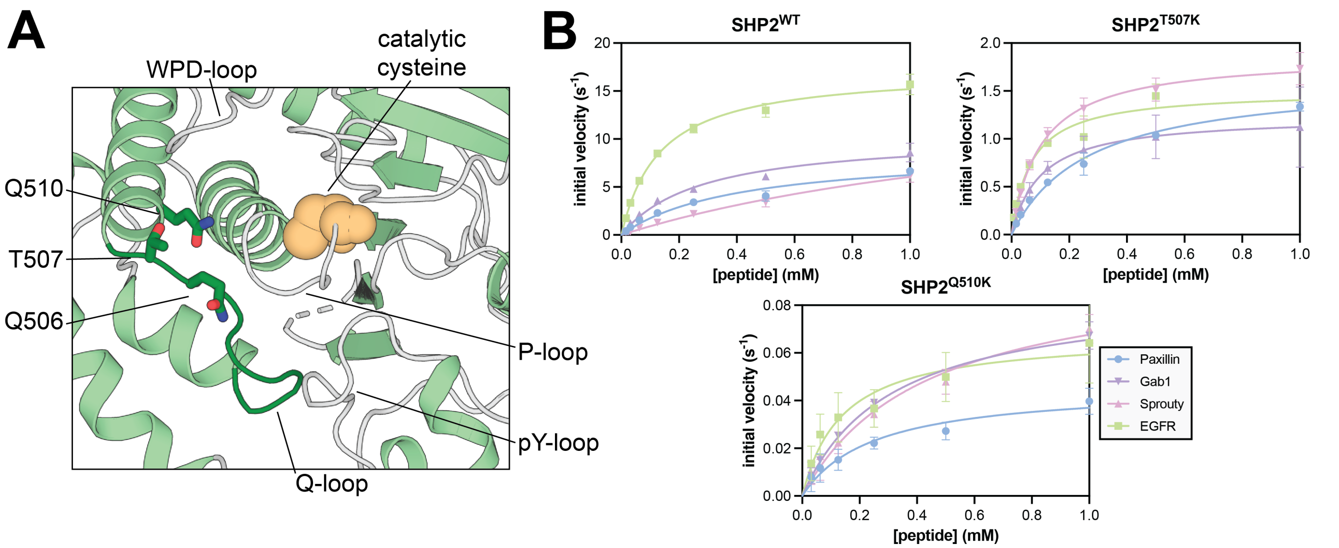


**Supplementary Figure 5. Michaelis-Menten analysis of PTP$^{T507K}$ and PTP$^{Q510K}$.** (**A**) Structure of SHP2 catalytic pocket. Catalytic cysteine and key catalytic loops are indicated (pY-loop: residues 276–282, WPD-loop: residues 420–429, P-loop: residues 458–465, and Q-loop: residues 501–507. Q510, T507, and Q506 are indicated as sticks. (**B**) Michaelis-Menten curves for PTP$^{WT}$, PTP$^{T507K}$ and PTP$^{Q510K}$ with four different peptides. Data are derived from are averages of 3 or more independent peptide and protein dilutions and measurements.

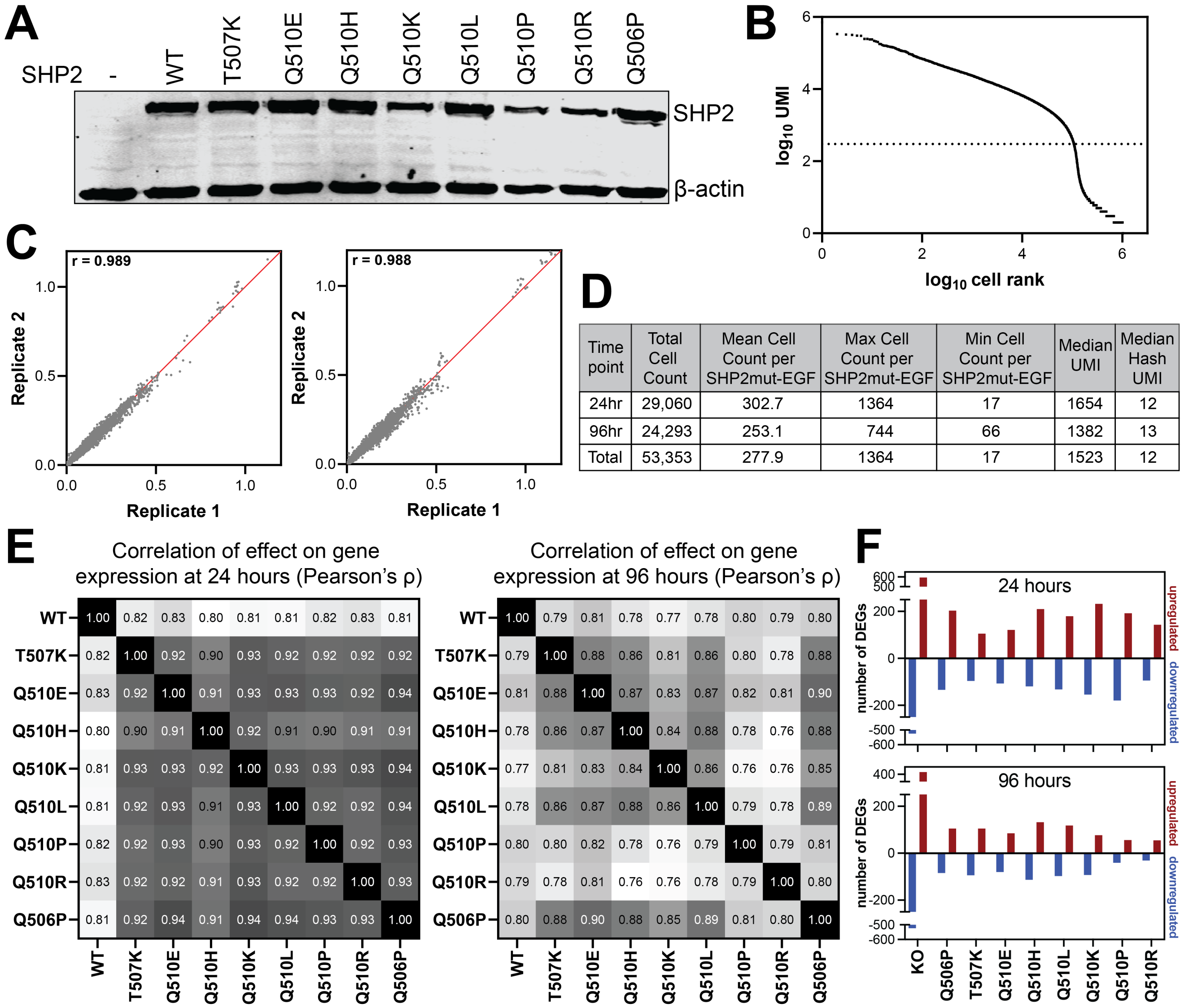


| Time point | Total Cell Count | Mean Cell Count per SHP2mut-EGF | Max Cell Count per SHP2mut-EGF | Min Cell Count per SHP2mut-EGF | Median UMI | Median Hash UMI |
|---|---|---|---|---|---|---|
| 24hr | 29,060 | 302.7 | 1364 | 17 | 1654 | 12 |
| 96hr | 24,293 | 253.1 | 744 | 66 | 1382 | 13 |
| Total | 53,353 | 277.9 | 1364 | 17 | 1523 | 12 |

**Supplementary Figure 6. Screening of Q-loop mutants.** (**A**) Western blot shows expression of $SHP2^{WT}$ and mutants included in the Q510 follow-up screen, compared to parental $SHP2^{KO}$ cell line. (**B**) Knee plot as UMI vs cell rank with a UMI cut-off of 300. (**C**) Correlation (pearson's r) between two independent replicates at 24 hours (*left*) and 96 hours (*right*). (**D**) Table of experimental summary metrics. (**E**) Correlation of β coefficients of SHP2 variants in Q510 screen at 24 hours (*left*) and 96 hours (*right*). (**F**) Number of differentially expressed genes (normalized effect size <-0.25 or >0.25, false discovery rate <0.05) for each SHP2 variant against $SHP2^{WT}$ at 24 hours (*top*) and 96 hours (bottom).

### Methods

#### Cell culture

Cells were cultured in a 37 °C tissue culture incubator with 5% $CO_2$. Cells were discarded by passage 25 and tested for mycoplasma every 3 months. HEK 293 cells were grown in Dulbecco's Modified Eagle Medium (DMEM) with 10% Fetal Bovine Serum (FBS) and 1% penicillin/streptomycin. HEK 293 SHP2 knock-out cells were grown in DMEM with 10% FBS. Human epidermal growth factor was purchased in lyophilized form (#E9644, Sigma) and reconstituted in 10 mM acetic acid.

#### Transfection and EGF stimulation

$10^6$ HEK 293 $SHP2^{KO}$ cells were seeded in a 6-well plate, and transfected 24 hours later with 2 µg DNA in 200 µL empty DMEM using 6 µL of polyethylenimine (PEI). Early the next morning, each well was trypsinized and reseeded into a 96-well plate to a final density of 10,000 cells per well. 12 hours after re-seeding, cells were checked for adherence and culture media was aspirated. Cells were serum-starved overnight in DMEM + 0.2%FBS. After 12 hours of serum-starvation, EGF was added to the cells in DMEM + 0.2% FBS to a final concentration of: 1000 ng/mL, 500 ng/mL, 250 ng/mL, 100 ng/mL, 50 ng/mL, 25 ng/mL, 12.5 ng/mL, 0 ng/mL (unstimulated). At 24 hours and 96 hours, cells were harvested and nuclei were hashed and frozen at -80 °C until further processing.

#### Nuclei preparation and hashing

For both screens, cells were trypsinized and moved to V-bottom plates at 24- and 96-hours post-EGF exposure. Upon washing with ice-cold 1X Phosphate Buffered Saline (PBS), cells were lysed with EZ Lysis Buffer (Sigma) supplemented with 1% diethyl pyrocarbonate (Sigma), 0.1% Superase RNase Inhibitor (Thermo Fisher), and 500 fmol of hashing oligo. After lysis, nuclei were fixed with the addition of 1.25% formaldehyde in 1.25X PBS (to a final well concentration of 1% and 1X, respectively) and incubated on ice for 10 minutes. Nuclei were pooled into a plastic reservoir and moved into a 50mL conical for centrifugation at 650*g* for 5 minutes at 4°C. Supernatant was removed from the nuclei pellet, and nuclei were washed once with nuclei suspension buffer (NSB; 10 mM Tris-HCl, pH 7.4, 10 mM NaCl, 3 mM MgCl2, 1% Superase RNA Inhibitor (Thermo Fisher), 1% 0.2mg/mL Ultrapure BSA (New England Biosciences)). Nuclei were resuspended in NSB, slow-frozen in 10% DMSO, and stored at -80°C until sci-RNA-seq library preparation.

#### Library preparation and sequencing

Hashed nuclei were thawed and subjected to 3-level combinatorial indexing protocols adapted from previous methods[1–4]. Nuclei were spun, resuspended in NSB, and sonicated at low power for 12s (Bioruptor). Upon counting, 21µL nuclei were moved to 96-well low adhesion PCR plates with 2µL 10mM dNTP, 2µL 100 µM indexed oligo-shortdT primers, 2µL 100 µM indexed random hexamer primers, and 14µL of a reverse transcription master mix consisting of 14.29% 100mM DTT, 14.29% 100mM RNaseOUT Ribonuclease Inhibitor, 57.14% 5X SuperScript IV First-Strand Buffer, and 14.29% SuperScript IV Reverse Transcriptase. Reverse transcription was carried out with an increasing temperature gradient. Post-reverse transcription, nuclei were pooled and distributed as 10µL into a 96-well plate(s) for ligation steps. Briefly, 8µL of indexed ligation primers were added to each well, along with a 4.8µL 3:2 master mix of T4 ligase buffer:T4 ligase (New England Biosciences, NEB) and 9.4µL of nuclei buffer with BSA (NBB; 10 mM Tris-HCl, pH 7.4, 10 mM NaCl, 3 mM MgCl2, 1% 0.2mg/mL Ultrapure BSA). Ligation was carried out at 25°C for 1 hour. Resulting nuclei were pooled, washed with NBB, and distributed as 1500 nuclei in 5µL NBB per well, where some plates were stored for future processing. Next, 5µL of a second strand synthesis mix consisting of 60% elution buffer (Qiagen), 27% second strand synthesis buffer (NEB), and 13% second strand synthesis enzyme mix (NEB) was added, and second strand synthesis was carried out at 16°C for 3 hours. Post- second strand synthesis, tagmentation was performed at 55°C for 5 min after the addition of 1/50 µL of N7-adaptor loaded Tn5 and subsequent quenching with DNA binding buffer (Zymo) for 5 min at room temperature. Resulting dsDNA was purified using a 1X SPRIbead clean-up within the 96-well plate. The dsDNA was eluted in buffer EB then moved to a clean 96-well PCR plate. To the 16µL of eluted product, 2µL P5 PCR primer and 2µL P7 PCR primer were added to wells in an indexed well-specific combination. Further, 20µL 2X NEBnext PCR master mix (NEB) was added, and PCR to add the adaptors was carried out. The final PCR product was pooled and subjected to a 0.7X SPRIbead cleanup for library cDNA purification and 1X cleanup for hash fraction purification. Library concentrations were determined by Qubit (Invitrogen) and were visualized by TapeStation DNA D1000. The resulting libraries were sequenced on the Element Biosciences AVITI for the initial SHP2 mutant screen according to the manufacturer's instructions and on the Illumina NovaSeq XPlus (Novogene) for the follow-up SHP2 Q510 screen.

### Data preprocessing and generation of count matrix

Raw base call files were obtained from Illumina BaseSpace or AVITI storage and were used to generate fastq files using bcl2fastq v2.20.0.422 or bases2fastq version 1.5.0.962525890, respective of sequencing platform. A custom data processing pipeline, adapted from previous publications[5], was used to process fastq data into a single-cell count matrix. First, reverse transcription and ligation barcodes were assigned to reads with a mismatch allowance of 1 base pair, and reads assigned to oligo-shortdT primers were separated from those assigned to random hexamer primers. After index assignment, polyA sequences were trimmed using TrimGalore version 0.6.10 and CutAdapt version 2.6. Upon polyA trimming, reads were aligned to human GRCh38 using the STAR aligner version 2.7.9a. Aligned reads were filtered for quality and duplicates and were assigned to genes using bedtools version 2.26.0, as described previously. The resulting unique read assignments from both primers were combined and collapsed by cell and gene, and a *celldataset* (CDS) object was generated using the raw sparse count matrix, cell annotations, and gene annotations with the R package *monocle3*. Cell barcodes were determined to be cells upon filtering the CDS by a cutoff of 300 unique molecular identifiers (UMIs), determined visually with the kneeplot of cell rank by UMI count. Lastly, doublets were detected with *scrublet* and were filtered based on the doublet score distribution.

In parallel, hash assignments were determined from demultiplexed untrimmed oligo-shortdT reads as described previously[5,6]. Briefly, hash barcodes were assigned to reads with a mismatch allowance of 1 base pair and if the read was adjacent to repeated A sequences, corresponding with hash sequence design. Duplicate hash reads were filtered by UMI and were collapsed into hash assignment counts by cell. Hashes were assigned to cells by two criteria: (1) a cell having ≥5 hash UMIs and (2) a ratio of the cell's top hash UMI to second best hash UMI of 2.5. The *monocle3* package was used to manipulate, batch align and visualize the resulting data.

### SHP2-mutant specific differential gene expression test

SHP2 variant was annotated in the object *colData* as a factor in order to designate either KO or WT as the reference comparison group. Differentially expressed genes (DEGs) were calculated by fitting expression to a quasi-poisson regression model of a gene within each timepoint as a function of SHP2 variant, EGF exposure concentration, and technical replicate using the *monocle3* R package *fit_models* function. The tests were limited to genes expressed in at least 1% of all cells in the experiment. P-values for each DEG test were false-discovery rate (FDR) corrected (specifically, Benjamini-Hochberg correction for multiple hypotheses). Significant SHP2 variant DEGs were defined as genes where the SHP2 variant coefficient was FDR < 0.01 and a normalized effect magnitude (β-coefficient) of > 0.05, unless otherwise indicated. The formula for obtaining SHP2-mutant specific normalized effects was used as follows:

$$Expression_{timepoint} \sim \beta_{intercept} + \beta_{SHP2\ variant}x_{SHP2\ variant} + \beta_{[EGF]}x_{[EGF]} + \beta_{replicate}x_{replicate} + \beta_{library\ size}x_{library\ size}$$

### EGF-dose specific differential gene expression test

For each SHP2 variant, differentially expressed genes were calculated by fitting expression to a quasi-poisson regression model of a gene as a function of EGF exposure concentration ($\log_{10}$ transformed with a pseudo-count of 0.1), timepoint, and technical replicate using the *monocle3* R package *fit_models* function. The tests were limited to genes expressed in at least 1% of all cells in the experiment. P-values for each DEG test were FDR corrected (Benjamini-Hochberg correction for multiple hypotheses). Significant EGF DEGs for each variant were defined as genes where the EGF dose coefficient was FDR < 0.01 and a normalized effect magnitude (β-coefficient) of > 0.05, unless otherwise indicated. The formula for obtaining EGF-dose specific normalized effects for each mutant is as follows:

$$Expression_{SHP2\ variant} \sim \beta_{intercept} + \beta_{[EGF]}x_{[EGF]} + \beta_{timepoint}x_{timepoint} + \beta_{replicate}x_{replicate} + \beta_{library\ size}x_{library\ size}$$

### Gene set enrichment analysis of SHP2$^{WT}$ and SHP2$^{KO}$ EGF-driven DEGs

The *fgsea* R package was used to assess the enrichment of gene sets in EGF-induced differentially expressed genes for SHP2$^{WT}$ and SHP2$^{KO}$. Briefly, for SHP2$^{WT}$ and SHP2$^{KO}$ separately, EGF-induced DEGs were ranked by descending effect ($\beta$-coefficient) and were tested for enrichment using the *fgsea* function against gene sets found in Hallmark pathways (h.all.v6.0.symbols) and Reactome pathways (c2.cp.reactome.v2024.1.Hs.symbols), downloaded from MSigDB. Myc-targets v1 was included to allow identification of broad effects, whereas Myc-targets v2 allows for detecting more direct Myc-driven effects. P-values for each test were FDR corrected (Benjamini-Hochberg correction for multiple hypotheses), and significantly enriched gene sets were designated as FDR < 0.05.

### Defining SHP2$^{WT}$ and SHP2$^{KO}$ EGF-driven gene modules

Gene modules were defined using the union of EGF-induced DEGs for SHP2$^{WT}$ and SHP2$^{KO}$ (FDR < 0.05). Upon filtering the cds object to SHP2$^{WT}$ and SHP2$^{KO}$ cells and EGF-induced DEG union genes, gene modules were defined using the *monocle3* R package *find_gene_modules* function with a resolution of 1e-2. Gene modules were validated by visual inspection of silhouette plots. For each cell and gene module, expression of genes was aggregated, and the mean expression for each SHP2 variant and EGF exposure group was z-scored for visualization with the R package *ComplexHeatmap*.

### SHP2 variant pair-wise correlation coefficient visualization

SHP2 variant pair-wise Pearson correlation coefficients were calculated using the normalized effects ($\beta$-coefficients) of the union of differentially expressed genes across SHP2 variants (FDR < 0.01 and abs(normalized effect) > 0.05). Unsupervised hierarchical clustering (ward.D2) and visualization of the correlation coefficients was performed using *ComplexHeatmap*.

### MrVI model training

MrVI was downloaded and imported from the publicly available *scvi-tools*. For the first SHP2 mutant screen, feature selection consisted of taking the union of the top 300 highly variable genes (determined with *scanpy*) for each timepoint_SHP2 variant subset of cells, for a total of 3318 genes.

We then trained a MrVI model on all 24-hour post-EGF exposure cells with the sample key defined as unique SHP2 variant, EGF concentration combination and the batch key defined as replicate. For all trained models, we used the recommended default model arguments: *n_latent=30; n_latent_u=10; qz_nn_flavor="attention"; px_nn_flavor="attention"; use_map (qz_kwargs)=True; stop_gradients (qz_kwargs)=False; stop_gradients_mlp (qz_kwargs)=True; dropout_rate (qz_kwargs)=0.03; stop_gradients (px_kwargs)=False; stop_gradients_mlp (px_kwargs)=True; h_activation (px_kwargs)="nn.softmax"; low_dim_batch (px_kwargs)=True; dropout_rate (px_kwargs)=0.03; learn_z_u_prior_scale=False; z_u_prior=True;* and *u_prior_mixture=False.* We used the following training arguments in tandem with the recommended default model arguments: *max_epochs=400; batch_size=256; early_stopping=True; early_stopping_patience=15; check_val_every_n_epoch=1; train_size=0.9; lr (pl_kwargs)=2e-3; n_epochs_kl_warmup (plan_kwargs)=20; max_norm (plan_kwargs)=40; eps (plan_kwargs)=1e-8;* and *weight_decay (plan_kwargs)=1e-8*.

For the follow-up Q510 screen, feature selection (2358 genes) and model training were conducted as with the preliminary screen. Separate models were trained upon filtering cells to respective EGF groups (No EGF, Low, High) in order to increase resolution to capture differences between mutants within respective EGF exposure groups.

### MrVI visualization and SHP2 variant cluster proportion

To visualize the output of the MrVI models, we generated UMAPs for the *U* and *Z* latent spaces. Specifically, the cell by *Z* latent dimensions were exported and added to the *cds* object in R where a UMAP embedding was generated and visualized using *monocle3 reduce_dimensions* and *plot_cells* functions. For downstream analysis, SHP2$^{KO}$ and Mock-transfected cells were filtered out to emphasize differences between remaining variants. Leiden community detection (*cluster_cells*) was performed at a resolution of 1e-3 and robustness of clustering was determined with silhouette plots. Counts of cells per cluster were used to determine cluster proportions for each SHP2 variant.

The functional relationship between the sample-unaware *U-space* and the sample-aware *Z-space* in MrVI can be used to directly estimate single-cell resolution sample-sample distance. After model training, both single-cell and mean sample-sample counterfactual distances were obtained using the *get_local_sample_distances* function. Single-cell counterfactual distances were visualized as distributions, and mean sample-sample counterfactual distance matrices were visualized as heatmaps using *ComplexHeatmap*.

### Purification of full-length SHP2 and SHP2 PTP domains

Full-length SHP2 variants were cloned into a pET28-His-TEV plasmid. BL21(DE3) cells were transformed with the respective plasmids, and were grown in TB supplemented with 100 µg/mL kanamycin at 37 °C until cells reached an OD600 of 0.5. IPTG (1 mM) was added to induce protein expression, which was carried out at 18 °C overnight. Cells were centrifuged and subsequently resuspended in lysis buffer (50 mM Tris pH 7.5, 300 mM NaCl, 20 mM imidazole, 10% glycerol, and freshly added 2 mM β-mercaptoethanol). The cells were lysed using sonication (Fisherbrand Sonic Dismembrator), and spun down at 14,000 rpm for 45 minutes. The supernatant was applied to a 5 mL Ni-NTA column (Cytiva). The resin was washed with 10 column volumes lysis buffer and wash buffer (50 mM Tris pH 7.5, 50 mM NaCl, 20 mM imidazole, 10% glycerol, and freshly added 2 mM β-mercaptoethanol). The protein was eluted off the Ni-NTA column in elution buffer (50 mM Tris pH 7.5, 50 mM NaCl, 500 mM imidazole, 10% glycerol) and brought onto a 5mL HiTrap Q Anion exchange column (Cytiva). The column was washed using Anion A buffer (50 mM Tris pH 7.5, 50 mM NaCl, 1 mM TCEP). Protein elution off the column was induced through a salt gradient between Anion A buffer and Anion B buffer (50 mM Tris pH 7.5, 1 M NaCl, 1 mM TCEP). The eluted protein was cleaved at the His6-TEV tag by addition of 0.10 mg/mL of His6-tagged TEV protease at 4 °C overnight. This cleavage cocktail was flowed through a 2 mL Ni-NTA gravity column (ThermoFisher) to separate the cleaved protein from uncleaved protein and TEV protease. Finally, the cleaved protein was purified by size-exclusion chromatography on a Superdex 200 10/300 gel filtration column (Cytiva) equilibrated with SEC buffer (20 mM HEPES pH 7.5, 150 mM NaCl, and 10% glycerol). Pure fractions were pooled and concentrated, and flash frozen in liquid N2 for long-term storage at −80 °C.

### Differential scanning fluorimetry

Purified protein stocks were thawed and diluted in DSF buffer (20 mM HEPES pH 7.5, 50 mM NaCl, 0.4% DMSO). 19 µL of buffer was added to a MicroAmp Fast Optical 96-well Reaction plate (Applied Biosystems, # 4346906). 1 µL of 500x SYPRO Orange Protein Gel Stain (Thermo Fisher, catalog no. S-6650) was added to a final protein concentration of 10 µM and 25x SYPRO Orange. Melting curves were performed in an Applied Biosystems Step-One Plus RT-PCR thermocycler. Temperature measurements started at 15 ºC, and temperature was raised by 0.5 ºC every minute with continuous measurements of fluorescence (excitation: 472 nm; emission: 570 nm). Raw fluorescence values along with corresponding temperatures were analyzed using DSFworld and Tm values were calculated using dRFU.

### DiFMUP basal activity measurements

Initial rate measurements for the SHP2-catalyzed dephosphorylation of 6,8-difluoro-4-methylumbelliferyl phosphate (DiFMUP) were conducted at 37 °C in DiFMUP buffer (60 mM HEPES pH 7.2, 150 mM NaCl, 1mM EDTA, 0.05% Tween-20). Reactions of 50 µL were set up in a black polystyrene flat bottom half area 96-well plate. A substrate concentration series of 31.25 µM, 62.5 µM, 125 µM, 250 µM, 500 µM, 1000 µM, 2000 µM and 4000 µM was used to determine $k_{cat}$ and $K_M$. Reactions were started by addition of appropriate amount of SHP2 wild-type and mutants (wild-type full-length: 2.5 nM; T507K full-length: 2.5 nM; Q510K full-length: 10 nM; Q510E: 2.5 nM; wild-type PTP: 1 nM; T507K PTP: 1 nM; Q510K PTP: 60 nM; Q510E PTP: 60 nM). Emitted fluorescence at 455nm was recorded every 25 seconds in a span of 50 minutes using a BioTek Synergy Neo2 multi-mode reader.

The linear part of the reaction progress curve was determined by visual inspection and fit to a line. Slopes were converted from absorbance or fluorescence units as a function of time to product formation as a function of time using standard curves measured with the reaction products (6,8-difluoro-7-hydroxy-4-methylcoumarin). Finally, these rates were corrected for enzyme concentration by dividing the values the concentration of enzyme used in the experiment to yield $V_0$ / [enzyme] in units of ($s^{-1}$). These corrected rates were plotted as a function of substrate concentration and fit to the Michaelis-Menten equation using non-linear regression to determine catalytic parameters. Experiments were generally repeated at least three times, and the average and standard deviation of all individual replicates are reported.

### Phosphatase activity cassay against phosphopeptide substrates

Initial rate measurements of the dephosphorylation of phosphopeptide substrates were conducted at 37 °C in freshly made reaction buffer (10 mM HEPES pH 7.5, 150 mM NaCl, 1mM TCEP, and 10% glycerol) using the EnzCheck Phosphate Assay Kit (ThermoFisher) according to the manufacturer's intstructions. Reactions of 50 µL were set up in a clear polystyrene flat bottom half area 96-well plate. A substrate concentration series of 0-1000 µM was used to determine $k_{cat}$ and $K_M$. Reactions were started by addition of 4000 nM SHP2. Absorbance at 360 nm was recorded every 8 seconds in a span of 6 minutes using a BioTek Synergy Neo2 multi-mode reader.

In all cases, the linear region of the reaction progress curve was determined by visual inspection and fit to a line. These slopes were converted from absorbance units as a function of time to product formation as a function of time using standard curves measured with inorganic phosphate. These rates were then corrected for enzyme concentration used in the experiment to yield $V_0$ / [enzyme] in units of ($s^{-1}$). This was plotted as a function of substrate concentration and fit to the Michaelis-Menten equation using non-linear regression to determine catalytic parameters. Experiments were repeated at least three times, and the average and standard deviation of all individual replicates are reported.

### Peptide synthesis and purification

All peptides were synthesized using 9-fluorenylmethoxycarbonyl (Fmoc) solid-phase peptide chemistry. All syntheses were carried out using the Liberty Blue automated microwave-assisted peptide synthesizer from CEM under nitrogen atmosphere, with standard manufacturer-recommended protocols. Peptides were synthesized on MBHA Rink amide resin (0.1 mmol scale). Each Nα-Fmoc-amino acid (6 eq, 0.2 M) was activated with diisopropylcarbodiimide (DIC, 1.0 M) and ethyl cyano(hydroxyamino)acetate (OxymaPure, 1.0 M) in N,N-dimethylformamide (DMF) prior to coupling. 0.4 molar-equivalents (0.4 M) of diisopropylethylamine (DIEA) was added to the OxymaPure solution, following the CarboMAX method published by CEM. Each coupling cycle was done at 75°C for 15 s followed by 90°C for 110 s. Deprotection of the Fmoc group was performed in 20% (v/v) piperidine in DMF (75°C for 15 s then 90°C for 50 s). The resin was washed 4x with DMF following Fmoc deprotection and after Nα-Fmoc amino acid coupling. All peptides were acetylated at the N-terminus with 10% acetic anhydride/DMF and washed 4x with DMF after the acetylation reaction.

Following peptide synthesis, the resin was washed 3x with DMF, dichloromethane (DCM), and methanol (MeOH) and dried under reduced pressure overnight. The peptides were cleaved from resin and the side chains were simultaneously deprotected in 95% (v/v) trifluoroacetic acid (TFA), 2.5% (v/v) water, and 2.5% (v/v) triisopropylsilane (TIPS), in a ratio of 10 µL of cleavage cocktail per mg of resin. The cleavage-resin mixture was incubated at room temperature for 90 min, with agitation. The cleaved peptides were precipitated in cold diethyl ether, pelleted, and dried under air. The peptides were dissolved in 50% (v/v) acetonitrile/water solution and filtered from resin. The filtrate was freeze-dried for downstream purification.

The crude peptide mixture was purified using reverse-phase high performance liquid chromatography (RP-HPLC) on a preparatory C18 column (Waters, XBridge Peptide BEH C18 OBD Prep Column, 19x150mm, 5µm). Flow rate was maintained at 17 mL/min with solvents A (water, 0.1% (v/v) TFA) and B (acetonitrile, 0.1% (v/v) TFA). Peptides were generally purified over a 23 min linear gradient from 0-70% solvent B. Peptide purity was assessed using an analytical column (Agilent, ZORBAX 300 SB-C18, 4.6x150mm, 5µm) at a flow rate of 1 mL/min over a 0-90% solvent B gradient in 30 min. All peptides were determined to be ≥95% pure by peak integration. The identities of the peptides were confirmed using mass spectrometry (Waters Xevo G2-XS QTOF). Pure peptides were lyophilized and redissolved in Tris buffer (100 mM, pH 8.0) as needed for experiments.
